# Targeting tumor antioxidant pathways with novel GSTP1/M2 inhibitors for cancer treatment

**DOI:** 10.1101/2025.10.20.683468

**Authors:** Greeshma P. Kumpati, Conor T. Ronayne, Joseph L. Johnson, Zachary S. Garder, Anuj K. Singh, Ananth Iyer, Venkatram R. Mereddy

## Abstract

Glutathione transferase (GSTP1 and GSTM2) are tractable targets for anticancer drug development. In this work, a series of 6-(7-nitro-2,1,3-benzoxadiazol-4-ylthio)hexanol (NBDHEX) based analogues were designed, synthesized, and evaluated both theoretically and experimentally as GSTP1 and GSTM2 inhibitors. Among the synthesized compounds, **3h** showed selective inhibition toward GSTP1 while **5b** showed selective inhibition of GSTM2. Compounds **5b** and **5c** exhibited stronger potency while compound **3h** showed slightly lower potency against the tested cancer cells than its parent molecule NBDHEX. Comprehensive biological studies were conducted on the effect of **3h, 5b** and **5c** towards breast cancer MDA-MB-231 and pancreatic MiaPaCa-2 cell lines revealed that **3h, 5b** and **5c** could activate JNK pathway and induce cell apoptosis. Furthermore *in vivo* experiments using NSG mice demonstrated that **5b** significantly reduced tumor growth when administered in combination with gemcitabine, effectively overcoming gemcitabine resistance in the MiaPaCa-2 cell model through targeted inhibition of GSTM2. These findings suggests that, **5b** could become a promising candidate for further development as a potential antitumor agent in cancer therapy.

## INTRODUCTION

Glutathione S-transferase (GSTs) are predominantly known phase II metabolizing enzymes that are potentially involved in emergence and development of various types of cancers.^1^ GSTs belong to a super gene family that includes cytoplasmic, mitochondrial, and microsome families.^2^ Among these, the cytoplasmic family contains alpha, mu, pi, sigma, theta, zeta, and omega subtypes, that are involved in various human diseases.^1–3^ The Pi class GSTP1 and mu class GSTM2 enzymes are frequently overexpressed in many cancers and contribute to chemoresistance by facilitating the elimination of toxic metabolites promoting tumor cell survival and inhibiting apoptosis.^4–5^ It was also reported that GSTM2 inhibition has been shown to enhance chemosensitivity and improve survival in pancreatic and other cancers.^6^

C-jun N-terminal kinase (JNK) is predominantly known to play an important role in a variety of physiological and pathological processes, including cell cycle, reproduction, apoptosis, and cell stress.^7^ Under normal cellular conditions, GSTP1 interacts with JNK forming adduct through protein-protein interactions that promote JNK inactivation, preventing cancer cells from undergoing apoptosis.^7,8^ Under cellular stress conditions, GSTP1 gets separated from JNK, where JNK can mediate the phosphorylation of the downstream c-jun protein and activate the cellular apoptosis.^8^ However, GSTP1 upregulation within the cells, can potentially block cellular apoptosis and contribute for drug resistance within cancer cells. In addition to that, the mu class enzyme GSTM2 binds to the N-terminal region of ASK1 and blocks the N-terminal dimerization of ASK1, leading to its inactivity, inhibiting the activation of ASK1-JNK/P38 signaling cascades. Thus, inhibition of GSTP1 and GSTM2 can prompt cell apoptosis.^9^

Over the years, significant progress has been made in the search for GST inhibitors. Among the GSTP1 and GSTM2 inhibitors developed to date include ethacrynic acid (EA) derivatives, glutathione analogues such as TLK117/119 and NBDHEX.^10–12^ NBDHEX (6-(7-nitro-2,1,3-benzoxadiazol-4-ylthio)hexanol) is a first-in-class inhibitor of GSTP1 and GSTM2 that effectively suppresses their enzymatic activity and disrupts critical protein-protein interactions with key MAPK proteins involved in cell survival.^12^ Despite its potent anticancer activity, NBDHEX exhibits off-target cytotoxicity and suboptimal pharmacokinetic properties, which limit its clinical utility.^12^ To address these challenges, we designed and synthesized two series of novel NBDHEX analogues with the goal of enhancing target binding affinity and metabolic stability while preserving the compound’s ability to interfere with oncogenic signaling complexes. The first series of analogues consisted of mono-NBD derivatives in which the hydrophobic alkyl chain bearing a terminal hydroxyl group[(CH_2_)_6_OH] was replaced with aromatic or heteroaromatic moieties such as benzene, pyridine, oxazole, thiazole, or imidazole rings. The second series comprised di-NBD derivatives containing two NBD units connected through linkers such as dithiothreitol (DTT) or polyethylene glycol (PEG). These structural modifications were designed to generate novel molecular entities with improved pharmacological and pharmaceutical properties.

## RESULTS AND DISCUSSION

### Synthesis of mono- and di-substituted analogues of nitrobenzoxadiazole derivatives as potential inhibitors of GSTP1 and GSTM2

Nitrobenzoxadiazole derivatives **(3a-3m)** were synthesized from commercially available 4-chloro-7-nitrobenzofurazan **(1)**, reacted with thiol containing functionalized molecules **(2a-2m)** in an equimolar mixture of ethanol and 1 M phosphate-buffered saline (PBS) at room temperature, with the reaction maintained at pH 7 outlined in **scheme 1.**^13^

After synthesizing the first generation of mono nitrobenzoxadiazole derivatives, we were motivated to extend the synthesis work of development of di-nitrobenzoxadiazole derivatives. The rationale was twofold with first, to enhance the cytotoxic potential of the compounds, and second to introduce more functionally versatile templates. In order to achieve this, we introduced dithiothreitol (DTT) containing derivatives **(5a & 5b)** and a PEG-containing derivative **5c (Figure 2)** to enhance both cytotoxicity and target selectivity. For the synthesis of derivatives **(5a-5c)**, a protocol similar to that used for the mono-substituted series was employed with 2 equivalents of 4-chloro-7-nitrobenzofurazan **(1)** with 1 equivalent of commercially available precursor molecules **(4a-4c)** as outlined in **scheme 2**.

**Figure 1.**
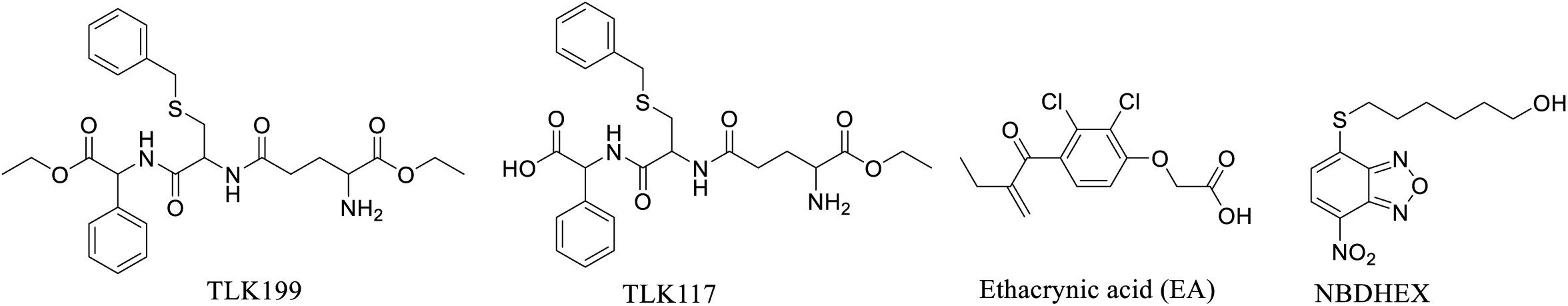
Structures of well-known GSTs inhibitors.

**Figure 2.**
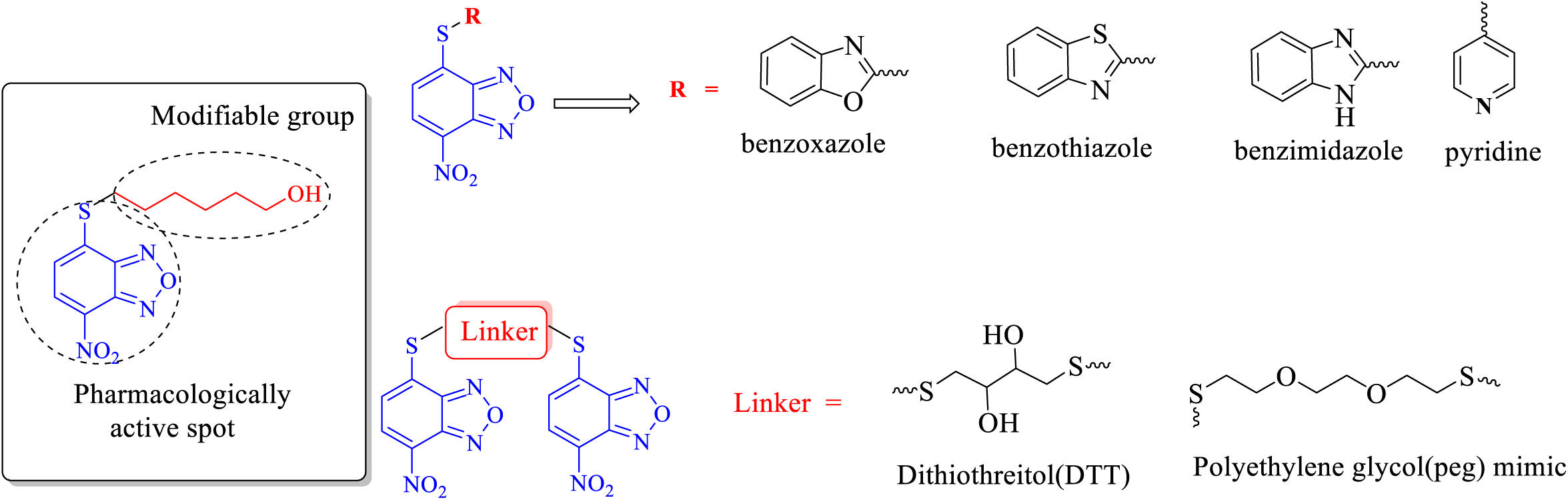
Strategy for the target NBD-based analogues.

### In vitro cancer cell proliferation inhibition assays illustrate that di-NBDHEX 5b and 5c exhibit potent anticancer properties

All the synthesized compounds **(3a-3m** and **5a-5c)** were evaluated for their potential *in vitro* anti-proliferative activity using the standard MTT colorimetric assay against a panel of four different cancer cell lines including human triple negative breast cancer cell line MDA-MB-231, human pancreatic cancer cell line MiaPaCa-2, metastatic murine breast cancer cell line 4T1 and non-metastatic murine breast cancer cell line 67NR respectively. These cell lines were selected based on their glutathione S-transferase (GST) expression profiles: MDA-MB-231 exhibits dual expression of GSTP1 and GSTM2, MiaPaCa-2 predominantly expresses GSTP1, and 4T1 and 67NR cells primarily express GSTM2 (**Figure S1**).

The benzoxazole, benzothiazole, and benzimidazole derivatives (**3b-3f**) showed reduced potency relative to the parent NBDHEX **(3a)**, however, they demonstrated greater selectivity for 67NR and MiaPaCa-2 (**Table 1**). In contrast, pyridine derivatives **(3g-3k)** improved overall activity, specifically with molecule **(3h)** emerging as the most potent analogue across the panel of the cell lines within the mono-nitrobenzoxadiazole conjugate series. Among the di-nitrobenzoxadiazole conjugate analogues, compound **(5a)** showed weakest activity, whereas compound **(5b)** displayed the strongest potency across all tested cell lines (**Table 1**). Notably, PEG-linked analogue **(5c)** also retained high cytotoxicity activity, comparable to compound **(5b)**, suggesting that linker modification can preserve or enhance biological activity. It is worth noting that compounds **3h, 5b** and **5c** were the most potent derivatives possessing broad-spectrum cytotoxicity against four different cancer cell lines among all the target compounds, so they were subjected to further biological evaluation.

**Table 1.** The inhibition rate (%) of target compounds (3a-3m and 5a-5c) against the panel of cancer cell lines MDA-MB-231, MiaPaCa-2, 4T1 and 67NR at 50 calculated as IC50. Data represents the average ± SEM of three independent experiments (n=3).

| SI. No. | MDA-MB-231 | MiaPaCa-2 | 4T1 | 67NR |
| --- | --- | --- | --- | --- |
| Positive control<br>NBDHEX(3a). | 3.37 ± 0.15 | 1.89 ± 0.51 | 1.32 ± 0.22 | 1.09 ± 0.06 |
| 3b. | 11.40 ± 0.26 | 8.37 ± 0.62 | 11.68 ± 0.53 | 8.67 ± 0.96 |
| 3c. | 12.88 ± 0.5 | 3.47 ± 0.72 | 11.6 ± 0.76 | 5.8 ± 0.15 |
| 3d. | 41.09 ± 0.56 | 19.04 ± 1.42 | 51.68 ± 0.92 | 14.6 ± 2.1 |
| 3e. | 11.18 ± 0.34 | 5.53 ± 0.22 | 11.09 ± 0.44 | 9.9 ± 1.3 |
| 3f. | 19.57 ± 2.30 | 4.41 ± 1.02 | 8.42 ± 0.63 | 7.03 ± 0.98 |
| 3g. | 12.20 ± 0.93 | 6.57 ± 0.81 | 8.53 ± 0.37 | 5.71 ± 1.3 |
| 3h. | 7.50 ± 0.26 | 3.49 ± 0.42 | 6.35 ± 1.45 | 4.52 ± 0.11 |
| 3i. | 10.35 ± 1.75 | 3.17 ± 0.61 | 8.10 ± 1.23 | 5.83 ± 1.85 |
| 3j. | 10.45 ± 0.54 | 10.22 ± 1.62 | 6.74 ± 0.93 | 5.91 ± 2.30 |
| 3k. | 11.91 ± 0.31 | 7.37 ± 1.08 | 5.58 ± 1.44 | 4.81 ± 1.31 |
| 3l. | 10.32 ± 1.33 | 4.82 ± 0.63 | 6.47 ± 0.81 | 5.81 ± 1.63 |
| 3m. | 14.16 ± 1.83 | 10.67 ± 0.98 | 8.41 ± 1.96 | 6.54 ± 1.23 |
| 5a. | 4.42 ± 1.45 | 4.98 ± 1.63 | 4.23 ± 0.96 | 3.41 ± 1.21 |
| 5b. | 1.11 ± 1.24 | 1.81 ± 0.93 | 1.90 ± 0.11 | 1.08 ± 0.23 |
| 5c. | 2.11 ± 0.57 | 1.78 ± 0.63 | 1.65± 0.25 | 0.93 ± 0.36 |

### Evaluation of 3h, 5b, and 5c as GSTP1 and GSTM2 Inhibitors via Docking and Enzyme Assays

Molecular docking and enzyme inhibition studies were performed to evaluate the binding and inhibitory potential of selected NBD-based analogues **3h, 5b** and **5c** against GST isoenzymes along with NBDHEX **3a**. Docking results revealed distinct isoform selectivity, with compound **3h** showing the strongest affinity for GSTP1 (E=-6.41 kcal/mol), comparable to NBDHEX **3a** (E = −6.22 kcal/mol), while compound **5b** bound less effectively (E = −5.77 kcal/mol) **Figure S2**. In contrast, compound **5b** exhibited markedly higher affinity toward GSTM2 (E = −10.85 kcal/mol) compared to NBDHEX **3a** (E = −7.50 kcal/mol) and compound **3h** (E = −5.42 kcal/mol), suggesting selective GSTM2 targeting **Figure S2**. Enzyme inhibition assays further confirmed that structural modifications introduced in the SAR studies significantly impacted GST isoform selectivity, with compounds **3h** and **5b** emerging as promising GSTM2 selective inhibitors, and compound **3h** as a potential GSTP1 selective scaffold as shown in **Table 2 & Figure S3**. Consistent with docking predictions, all three compounds were shown to bind to the H-site of GSTP1 and GSTM2, thereby blocking glutathione (GSH) conjugation. To determine the mode of inhibition, spectrophotometric assays were conducted in potassium phosphate buffer (pH 7.4) at 37 °C, monitoring GSH interactions with NBD-based compounds through UV-Visible spectra (200-600 nm) before and after acid treatment. These experiments demonstrated covalent conjugation of compounds **3h**, **5b**, and **5c** with GSH, confirming a covalent inhibition mechanism similar to NBDHEX **Figure S4**. Together, these findings establish compounds **5b** and **5c** as promising leads for selective GSTM2 inhibition and highlight compound **3a** as a basis for GSTP1 selective inhibitor design, providing valuable insights into GST isoenzyme-targeted drug development.

**Table 2:**
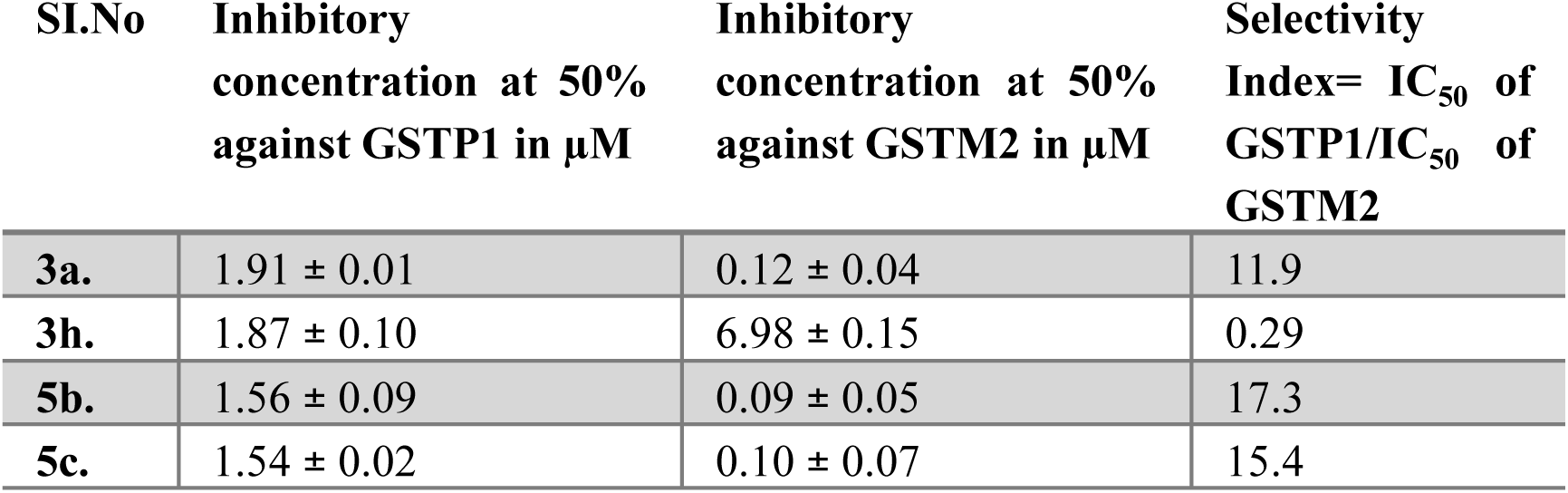
Inhibitory concentrations of the respective compounds against GSTP1 & GSTM2 along with their selectivity index values.

### Mechanism of Action for compounds 3a, 3h, 5b, and 5c involves the activation of JNK-mediated apoptosis

To investigate the mechanism of action of NBD-based analogues (**3h**, **5b**, and **5c**), we examined whether their cytotoxicity involved the JNK pathway, as previously reported for the parent compound NBDHEX (**3a**). Using the selective JNK inhibitor SP600125, we assessed whether the observed cell death was JNK-dependent or mediated through alternative signaling mechanisms.

To elucidate the mechanism of action of the synthesized NBDHEX (**3a)** and its analogues **3h, 5b** & **5c**, we examined their cytotoxic signaling pathways relative to the parent molecule NBDHEX (compound **3a**). Prior studies have shown that NBDHEX induces apoptosis primarily through activation of the JNK (c-Jun N-terminal kinase) pathway, a major component of the MAPK cascade.^8^**^&^**^12^ To determine whether our newly synthesized analogues act through similar mechanisms, we used selective inhibitor targeting JNK (SP600125) in MDA-MB-231 and MiaPaCa-2 cell lines (**Figure 3A**).

**Figure 3:**
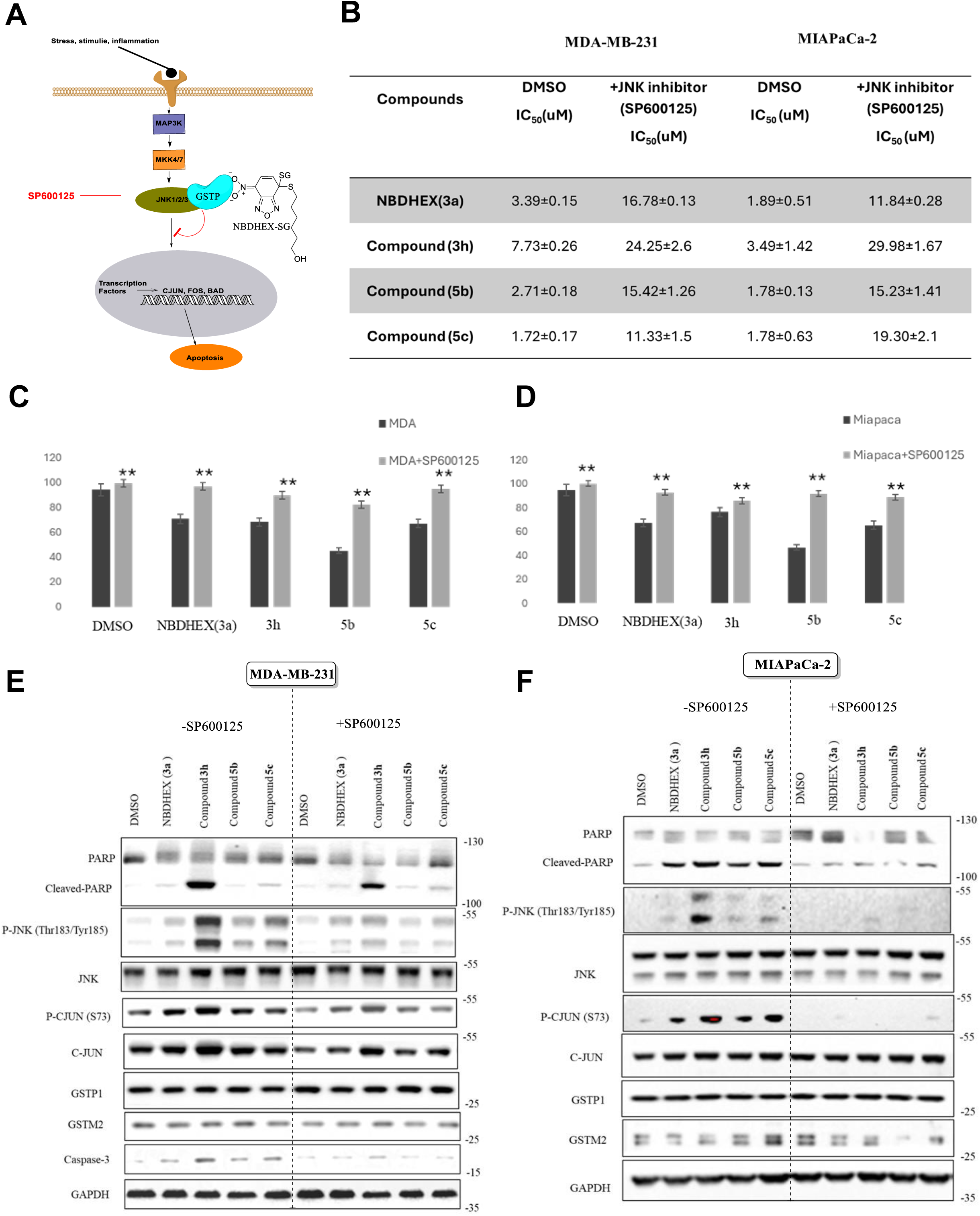
Compounds 3a, 3h, 5b & 5c activates the JNK pathway. a) MTT assay results to evaluate the cytotoxicity in the absence and presence of JNK inhibitor (SP600125) b) Model to illustrate the evaluation of **3a, 3h, 5b** & **5c** induced cytotoxicity through JNK-mediated apoptosis. c) SRB assay results to evaluate the cytotoxicity in the absence and presence of JNK inhibitor (SP600125). d) Western blot analysis illustrating the decrease in PARP, JNK, C-JUN activation in the presence of SP600125 upon treatment.

Co-treatment with the JNK inhibitor SP600125 markedly increased IC_50_ values for all compounds, confirming that JNK activation is essential for their cytotoxicity (**Figures 3B-D**, **S5** & **S6**). The most significant IC_50_ shifts were observed with compounds **5b** and **5c**, suggesting their strong dependence on JNK-mediated apoptosis highlighting JNK signaling as the primary cytotoxic mechanism of NBDHEX and its analogues (**Figure 3B**).

To further support these findings western blot analyses were conducted. Treatment with NBDHEX and its analogues significantly enhanced phosphorylation of JNK and its downstream target c-Jun, accompanied by cleavage of caspase-3 and PARP hallmarks of apoptosis. Among the analogues, compound **5b** elicited the strongest activation of p-JNK and p-c-Jun, correlating with its highest cytotoxic potency (**Figures 3E-F**). Importantly, co-treatment with SP600125 suppressed both phosphorylation and apoptotic marker cleavage, confirming that these effects are JNK-dependent (**Figure 3E-F**).

Together, these data demonstrate that NBDHEX and its newly designed analogues primarily induce apoptosis through activation of the JNK-c-Jun signaling axis. The dual NBD-core analogues, particularly compounds **5b** and **5c**, display enhanced potency likely due to stronger interactions with redox-sensitive targets and greater JNK activation. Overall, these findings establish a clear mechanistic basis for the cytotoxic activity of NBDHEX derivatives and highlight their potential as promising leads for the development of targeted anticancer agents that exploit JNK-mediated apoptotic signaling.

### GSTP1 and GSTM2 inhibitors sensitizes pancreatic cells towards gemcitabine treatment

Gemcitabine, an FDA-approved first-line drug for PDAC, often faces limited efficacy due to rapid chemoresistance.^14^ Studies have identified glutathione S-transferase mu 2 (GSTM2) as a key mediator of this resistance, particularly in MiaPaCa-2 pancreatic cancer cells.^9,13^ Gemcitabine treatment upregulates GSTM2 expression, while GSTM2 knockdown enhances apoptosis and reduces cell viability, indicating its role in resistance (**Figure 4A**).^13^ Based on these findings, we hypothesized that combining NBD analogues with gemcitabine would decrease the expression of GSTM2 and resensitize the MiaPaCa-2 cells towards gemcitabine treatment. *In vitro* studies in MIAPaCa-2 cells revealed that both compound **5b** and its combination with gemcitabine activated apoptotic pathways, as evidenced by increased cleavage of PARP, elevated P-JNK, P-c-JUN (**Figure 4B**). These results confirmed activation of the stress kinase pathway in the context of gemcitabine treated cells, consistent with the proposed mechanism of JNK-mediated apoptosis induced by NBD-based analogues.

**Figure 4:**
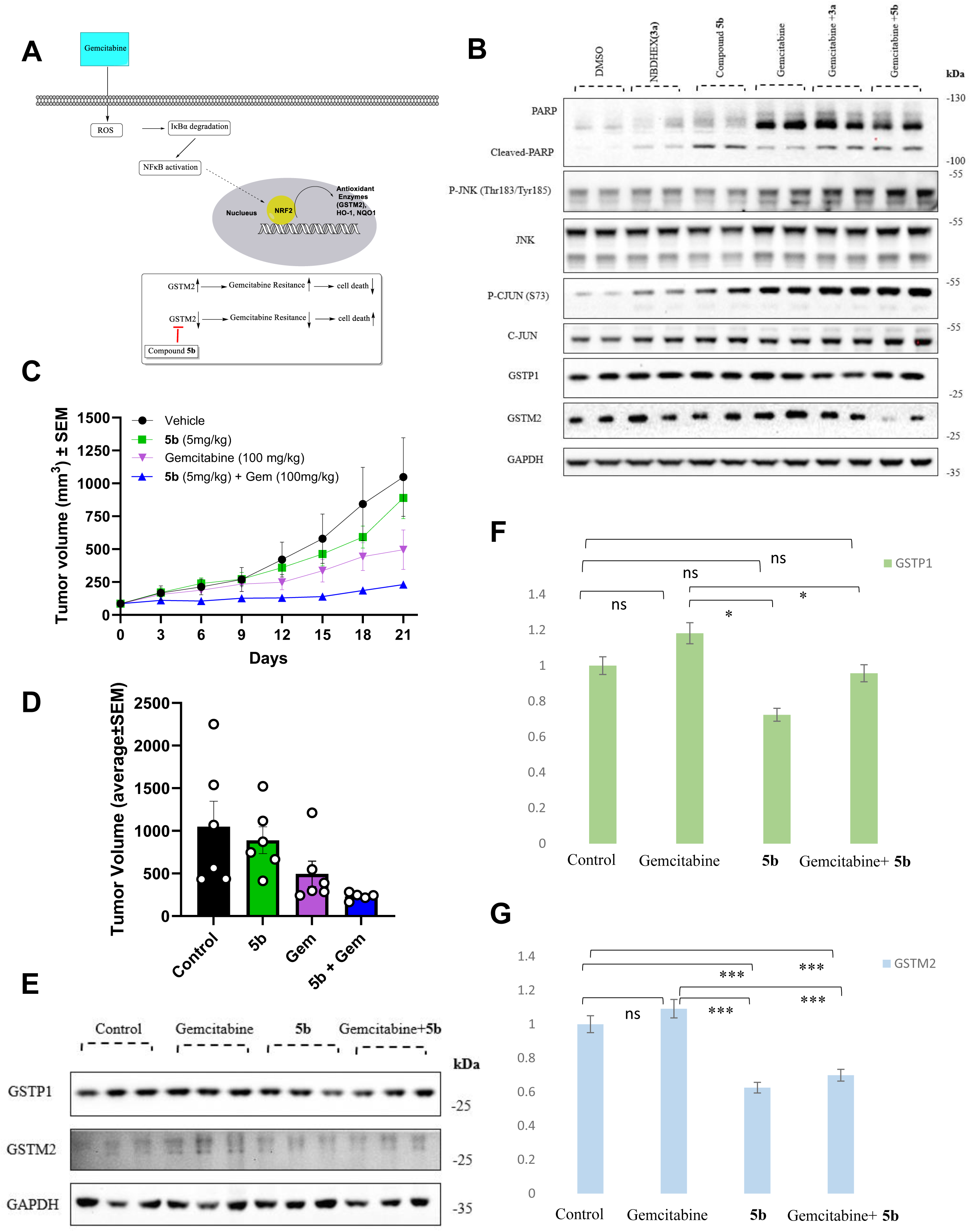
Antitumor efficacy and mechanistic analysis of compound 5b alone and in combination with gemcitabine. (a) Schematic representation illustrating the proposed mechanism of compound **5b** action. Compound **5b** inhibits GSTP1/GSTM2-mediated JNK suppression, leading to JNK activation and induction of apoptosis in combination with Gemcitabine.(b) Western blot analysis in vitro MiaPaCa-2 showing expression of apoptotic and signaling markers (PARP, Cleaved PARP, p-JNK, total JNK, p-c-JUN, and c-JUN) along with proteins of interest GSTP1, and GSTM2. (c) Tumor volume and rumor mass (d) analysis of MIA PaCa-2 xenograft-bearing mice treated with Vehicle, compound **5b** (5 mg/kg), gemcitabine (100 mg/kg), and their combination. Combination treatment significantly reduced tumor growth compared to single-agent treatments (mean ± SEM). (d) Western blot showing downregulation of GSTP1 and GSTM2 in tumor tissues upon combination therapy compared to control and single treatments. GAPDH served as the loading control. Doses were determined by carrying out an *in vivo* maximum tolerated dose study in healthy CD-1 mice (**supplementary Figure S7**) (e-f) Densitometric quantification of GSTP1 and GSTM2 normalized to GAPDH. Combination therapy significantly reduced GSTM2 expression (***p < 0.001) and modestly decreased GSTP1 (*p < 0.05), suggesting inhibition of GST-mediated detoxification and reversal of drug resistance.

To investigate the ability of GSTP1/M2 inhibition can resensitize tumors to gemcitabine, we carried out a tumor MIAPaCa-2 xenograft study (**Figure 4C-D**). MIAPaCa-2 tumors exhibit low levels of GSTM2 which can be induced by gemcitabine to cause treatment resistance. Hence, we envisioned that GST inhibition with compound **5b** might synergize with gemcitabine. Treatment with compound **5b** or gemcitabine alone did not result in substantial tumor growth inhibition by volume and mass (**Figure 4C-D**). However, combination therapy of **5b** with gemcitabine resulted in markedly suppressed tumor growth in MiaPaCa-2 xenograft models compared to control or single-agent gemcitabine (**Figure 4C-D**). Furthermore, combination treatment caused significant downregulation of GSTP1 and GSTM2 expression (**Figure 4E-G**), enzymes known to mediate detoxification and chemoresistance in gemcitabine treated tumors. The concurrent suppression of these GST isoforms and activation of JNK signaling suggests a synergistic mechanism by which compound **5b** enhances gemcitabine efficacy by impairing the tumor’s antioxidant defense and promoting apoptotic signaling. Overall, these findings demonstrate that compound **5b** potentiates gemcitabine’s antitumor effect via dual modulation of redox-regulating GST enzymes and activation of the JNK-mediated apoptotic cascade, offering a promising strategy to overcome chemoresistance in pancreatic cancer.

## CONCLUSIONS

GSTP1 and GSTM2 are tractable targets for anticancer drug development as these enzymes mediate cancer progression in advanced stage tumors and in the context of chemotherapy resistance. Here a series of NBDHEX-based drug candidates were designed, synthesized, and evaluated GSTP1 and GSTM2 inhibitors. *In vitro* GST-inhibition studies indicated that compound **3h** showed selective inhibition toward GSTP1 while **5b** showed selective inhibition of GSTM2. Compounds **5b** and **5c** exhibited stronger potency while compound **3h** showed slightly lower potency against the tested cancer cells than its parent molecule NBDHEX. Mechanistic studies were conducted on the effect of **3h, 5b** and **5c** towards breast cancer MDA-MB-231 and pancreatic MiaPaCa-2 cell lines revealed that **3h,5b** and **5c** could activate JNK pathway and induce cell apoptosis. Furthermore *in vivo* experiments using NSG mice demonstrated that **5b** significantly reduced tumor growth when administered in combination with gemcitabine, effectively synergizing and overcoming gemcitabine resistance in the MiaPaCa-2 xenograft model. This synergy was likely facilitated through inhibition of GSTM2. These findings position compound **5b** as a promising candidate for further development as a potential antitumor agent in cancer therapy.

## METHODS

### Materials and synthetic reagents

4-chloro-7-nitro-benzofurazan (NBD-Cl) and respective thiol (-SH) containing molecules such as were purchased from Ambeed (Arlington Heights, IL); The 1H- and 13C-NMR spectra were plotted on a Bruker Ascend™ 400 spectrometer. High-resolution mass spectra (HRMS) were recorded using a Bruker micrOTOF-Q III ESI mass spectrometer. Absorbance values were measured using BioTek Synergy 2 Multimode Microplate Reader (BioTek Instruments Inc., Winooski, VT, USA). All statistical analysis was carried out using GraphPad Prism 10 software (GraphPad Software, Boston, MA, USA).

### General Synthetic methods

Equal proportions of NBD-Cl **(1)** and respective thiols **(RSH)** were reacted for 6 hours in 20 mL of a 1:1 (v/v) mixture of ethanol and 0.1 M potassium phosphate buffer, pH 7.0 at 25 °C. The pH was constantly monitored with the help of pH stripes and kept neutral by suitable addition of 1 M KOH. Upon completion of the reaction, the excess RSH was removed by adding 1.5 mmol of 3-bromopyruvate to the reaction mixture. The solid reaction product, in 85-95% yields, were collected by filtration and washed twice with 15 mL of cold distilled water.

### Cell culture conditions

MDA-MB-231 cells (ATCC) grown in DMEM supplemented with 10% FBS and penicillin-streptomycin (50 μg/mL), MiaPaCa-2 cells (ATCC) grown in DMEM supplemented with 10% FBS, horse serum (2.5%), 4T1 cells (ATCC) and 67NR (University of Minnesota Duluth, Dr. Jon Holy) cultured in RPMI-1640 supplemented with 10% FBS and penicillin streptomycin (50 μg/mL). All cells were incubated at 37 °C and 5% CO_2_.

### MTT Cell Proliferation Assay

Cell cultures around 80-90% confluency were treated with trypsin followed by resuspension in the respective culture media to a concentration 5 × 10^4^ cells/ml. 100 µl of the 5 × 10^4^ cells/mL solution was added to a 95-well plate allowed to incubate at 37 °C, 5% CO2 for 24 hours. Compounds were then added to the cells in the plate and allowed to incubate for 72 hours. Upon incubation, 10 µL of MTT (5 mg/mL) was added to the 96-well plate and incubated for further 4 hours. After incubation, SDS (0.1 g/mL, 0.01 N HCl) was added and the 96-well plate was allowed to incubate for another 4 hours after which absorbance values were taken at 570 nm using a Biotek Synergy 2 plate reader. Absorbances of the respective compound treated wells were taken as percentages of the average untreated absorbances and plotted against log(concentration) using GraphPad Prism 10 to generate effective concentration values where 50% of the cells are not proliferating (IC_50_).

### Computational modeling

We evaluated the 3D binding interactions of the synthesized small-molecule compounds with GSTP1 and GSTM2 enzymes using autoDock Tools (ADT) 1.5.6. This computational docking analysis provided insights into the compounds’ binding affinities (kcal/mol) based on their lowest energy conformations. The chemical structures of the synthesized compounds were drawn using Chem Draw and converted into 3D structures via Open Babel, while the crystal structures of GSTP1 (PDB ID: 6GSS) and GSTM2 (PDB ID: 1XW5) were retrieved from the Protein Data Bank (PDB). All ligands were energy-minimized and converted into PDBQT format prior to docking. Grid and docking parameter files were generated before producing the docking log file, which was then analyzed to determine binding affinities and molecular interactions. A more negative binding energy value indicated stronger affinity between the compound and the target enzyme.

### GST kinetic Assay

The enzymatic activities of GSTP1 & GSTM2 (10 nM subunits) were measured spectrophotometrically at 340 nm (*ɛ* = 9600 M^-1^ cm^-1^) and 25 °C, by monitoring the rate of stoichiometric compositions of chloro-di-nitrobenzene (CDNB) conjugation with reduced glutathione (GSH) as a function of time. The reaction mixture contained 1 mM of CDNB and 1 mM of GSH in 1 mL of 0.1 M potassium phosphate buffer containing 0.1 mM EDTA and 0.1% (v/v) Triton X-100 maintained at pH 6.5. The inhibitory activity of the synthesized compounds was determined by recording the activity of GSTP1 and GSTM2 in the presence of various concentrations of the selected NBD derivative (0.05-50 μM). *K_i_* values for each compound were determined by fitting the data points to a hyperbolic saturation curve, and using GraphPad Prism 10 was used to generate effective concentration values where 50% of GSTP1 and GSTM2 activity was calculated (IC_50_).

### Spectrophotometric analysis

The spontaneous reactivity of the compounds with GSH was assessed by recording their UV visible spectra at 37^0^C. Each compound (50 µM) was dissolved in 0.1 M potassium phosphate buffer (pH 7.4) containing 0.1 mM EDTA in the presence of GSH (1 mM). The formation of GSH-compound complexes was evaluated by monitoring the emission spectra (λ_em = 250-600 nm) before and after GSH addition. To assess the stability of the resulting complexes, 6 M HCl was added after 6 hours of incubation, and the spectra were recorded again following an additional 6 hours.

### General tolerability of the NBDHEX and its derivatives in CD-1 mice

Female CD-1 mice (Charles River) were randomly divided into groups of six (n=6) after a week of mice acclimatization period, mice received intraperitoneal injections of respective compounds intraperitoneally starting at 5 mg/kg or vehicle control (10% DMSO, 10% PEG, 20% HS-15, and 60% sterile water) once for 4 days. Body weights were recorded and monitored on daily basis for behavior changes and/ or grooming pattern changes. Treatment was paused if body weight loss (BWL) exceeded 10%, and mice were euthanized if BWL surpassed 20%, following the approved protocol. The dosing was doubled every four days until 30 mg/kg, at which point the study was terminated due to signs of acute respiratory distress indicating compound toxicity. All animals were euthanized according to IACUC guidelines.

### Western Blot Analysis

Cells were seeded at a density of 150,000 cells/mL in 60 mm culture dishes and incubated overnight at 37 °C with 5% CO_2_. Following incubation, cells were treated with the indicated concentrations of test compounds for 24 hours. After treatment, cells were washed with 1× PBS and lysed using 100 μL of 1× RIPA buffer (Millipore Sigma, 20188) supplemented with phosphatase (50×, Sigma Aldrich) and protease (10×, Sigma Aldrich) inhibitor cocktails. The lysates were collected by scraping, sonicated for 10 seconds, and centrifuged at 13,000 × g for 5 minutes at 4 ^0^C. Supernatants were collected, and protein concentrations were determined using a BCA assay. Equal amounts of protein (20 μg) were resolved on NuPAGE 4-12% Bis-Tris gels (Thermo Fisher Scientific, NP00336BOX) alongside a pre-stained protein ladder (Thermo Scientific, 26619), using MOPS SDS running buffer (Thermo Fisher, NP000102), and transferred onto PVDF membranes.The membranes were probed with the following primary antibodies (1:1000 dilution): IRE1α, p-IRE1α, JNK, p-JNK, c-Jun, p-c-Jun, PARP, cleaved caspase-3, GSTP1, GSTM2, and GAPDH (Cell Signaling Technology, Thermo Fisher, and Abclonal). After incubation with HRP-conjugated secondary antibodies (1:10,000), protein bands were visualized using enhanced chemiluminescence substrates (SuperSignal West Femto Luminol/Enhancer and Stable Peroxide, Thermo Scientific).

### SRB Cell Proliferation Inhibition Assay

Cells were seeded at a density of 3 × 10⁶ cells per well in 6-well plates and incubated overnight at 37 ^0^C with 5% CO_2_. The following day, cells were treated with the desired concentrations of test compounds and incubated for 24 hours under the same conditions. After treatment, cells were washed three times with cold PBS and air-dried overnight in a non-CO₂ incubator to fix. Each well was then stained with 100 µL of 0.5% (w/v) Sulforhodamine B (SRB) prepared in 1% aqueous acetic acid and incubated at 37 ^0^C for 45 minutes. The wells were rinsed with 1% acetic acid to remove unbound dye and air-dried. Finally, the bound SRB dye was solubilized in 10 mM Tris buffer (pH 10), and absorbance was measured at 540 nm.

### Compound 5b Tumor efficacy study in a NSG mice

NSG mouse were administered with MiaPaCa-2 cancer cells at 500,000 cells/mouse into their flank as a 1:1 solution of media and Matrigel making up to the total volume of 0.1 mL/1000 μL. Tumors were grown under proper monitoring until they reached a volume of ∼100 mm^3^ and were then randomly sorted into groups (n = 6). Mice were then administered intraperitoneally with vehicle control (10% DMSO,10% Peg, 20% Tween 80 & 60% saline water) and **5b** at 5 mg/kg once daily for 14 days. Body weights were recorded daily and monitored for behavior and grooming patterns. A body weight loss (BWL) >10% indicated a halting of treatment until weight recovery. If a BWL >20% was observed, that mouse would be euthanized. Tumor volume (TV, mm^3^) was calculated as follows: TV = (a×b2)/2, a as the tumor length and b as the tumor width. The tumors were measured every 2-3 days using a caliper in two dimensions. Mice were euthanized at the end of the 10-day treatment period where tumors were then resected and weighed. The study was carried out under IACUC protocol 2106-39167A. Statistical analysis was carried out via unpaired t-test using GraphPad Prism 10 (*p<0.05, **p<0.01, ***p<0.001, ****p<0.001).

### Protein extraction from tumors

The ceramic mortar, pestle, spatula, and Eppendorf tubes were pre-chilled on dry ice for at least 20 minutes before use. A portion of the snap-frozen tumor tissue was placed in the cooled mortar and ground into a fine powder using the pre-chilled pestle. Approximately 10-20 mg of the powdered tissue was transferred into a labeled tube containing 250 µL of ice-cold RIPA buffer (1×, Millipore Sigma, 20188) supplemented with phosphatase inhibitor cocktail (1×, Sigma-Aldrich, 50× easy pack) and protease inhibitor cocktail (1×, Sigma-Aldrich, 10× easy pack). The tissue suspension was sonicated for 5-10 seconds to achieve complete homogenization and then centrifuged at maximum speed for 10 minutes at 4 ^0^C. The resulting supernatant was collected and used for BCA protein quantification and Western blot analysis.

### Ethical Considerations

The general tolerability study in CD-1 mice was carried out in accordance with the guidelines and regulations of the University of Minnesota’s Institutional Animal Care and Use Committee (IACUC) under protocol [2211-40546A]. All experimental procedures were reviewed, approved, and adhered to institutional standards. Similarly, the 67NR tumor syngraft experiments were performed following approval from the University of Minnesota IACUC [2106-39167A]. This work was conducted in full compliance with all relevant guidelines and regulations, with protocols approved by the University of Minnesota.

### Statistics

Statistical analysis was conducted using GraphPad Prism 10 software (GraphPad Software, Boston, MA, USA). In vitro assays were performed independently in triplicate with DMSO as a control. Data was analyzed using Student’s t-test and ANOVA and log transformation was used when appropriate. P values <0.05 were considered statistically significant.

## SCHEME CAPTIONS

**Scheme 1.**
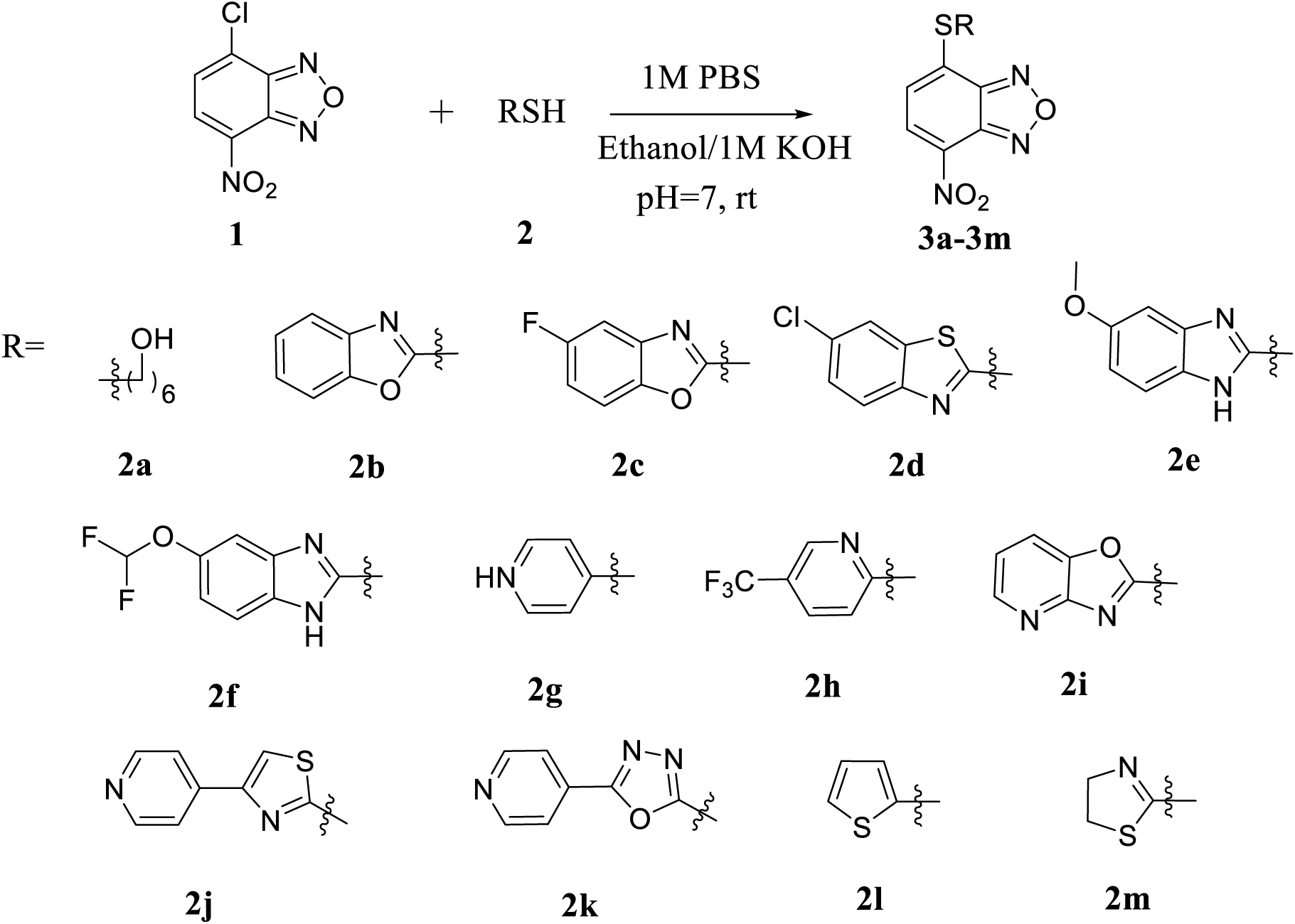
Synthesis of mono-nitrobenzoxadiazole unit containing derivatives.

**Scheme 2.**
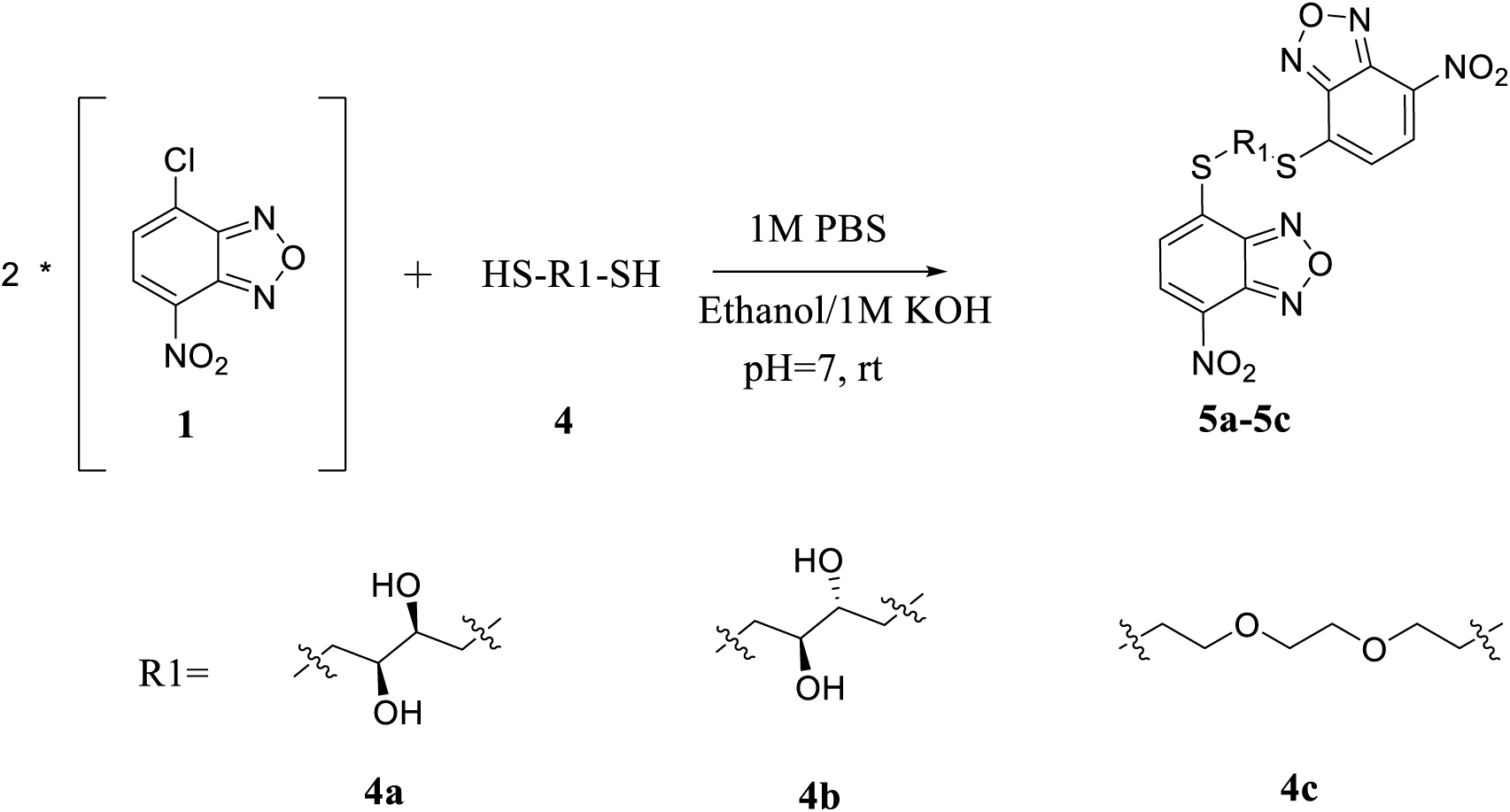
Synthesis of di-nitrobenzoxadiazole unit containing derivatives.

## SUPPLEMENTARY FIGURE CAPTIONS

**Figure S1.**
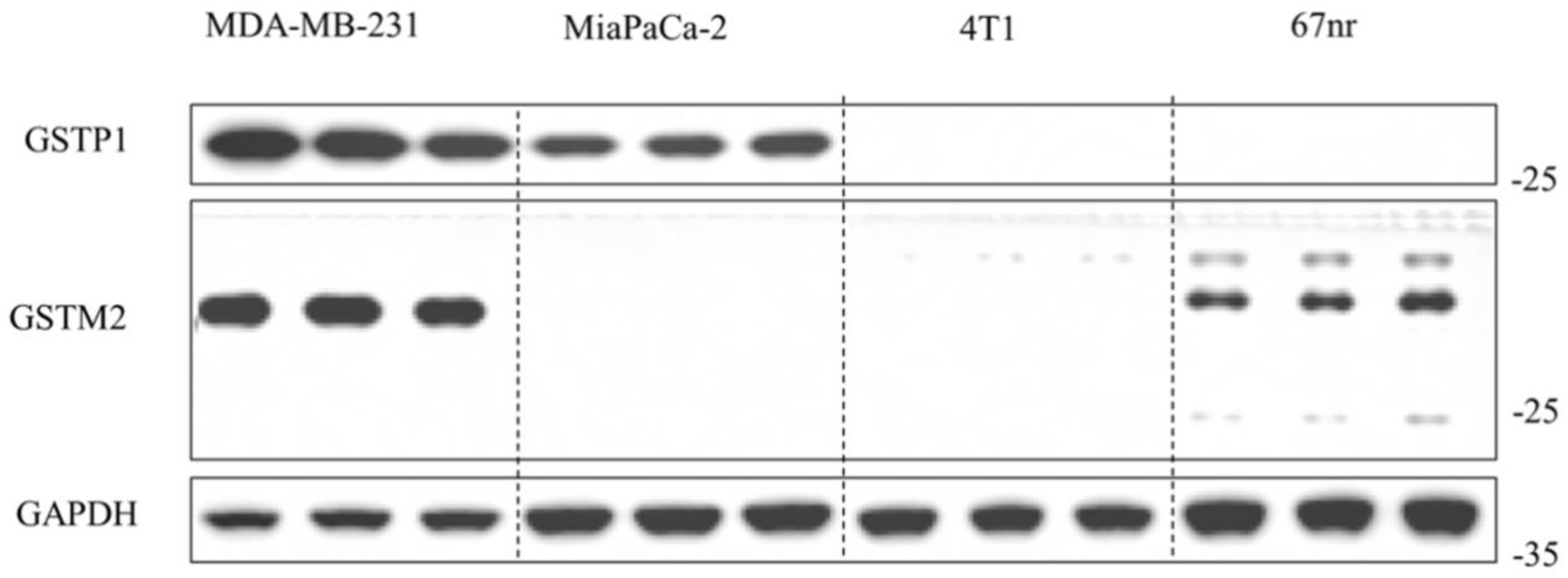
Representative Western blot showing the expression levels of GSTP1 and GSTM2 across four different cell lines: MDA-MB-231, MiaPaCa-2, 4T1, and 67NR. Proteins were collected from each cell line in triplicate, normalized to total protein using bicinchoninic acid (BCA), separated proteins by SDS-PAGE, and transferred to PVDF membranes. Membranes were probed with antibodies specific to GSTP1, GSTM2, and the loading control GAPDH.

**Figure S2.**
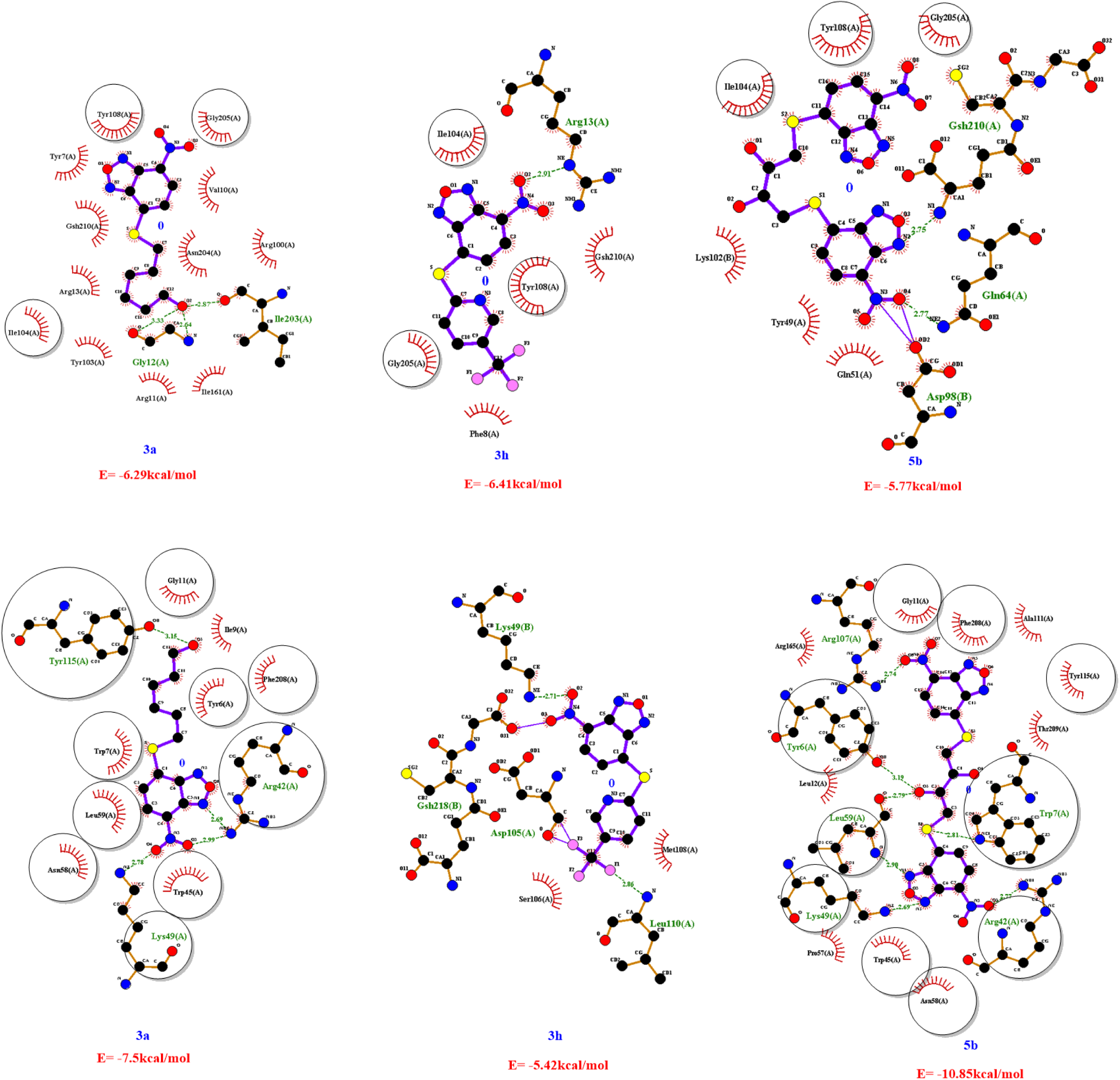
2D interaction diagrams of compounds **3a**, **3h**, and **5b** bound to the active site of GSTP1(A) and GSTM2(B), along with their corresponding binding energies. Hydrogen bonds were depicted as green dashed lines, while hydrophobic interactions were shown as red semicircles. Residues encircled in the figure represent the common amino acids within the active site of GSTP1 and GSTM2 respectively.

**Figure S3.**
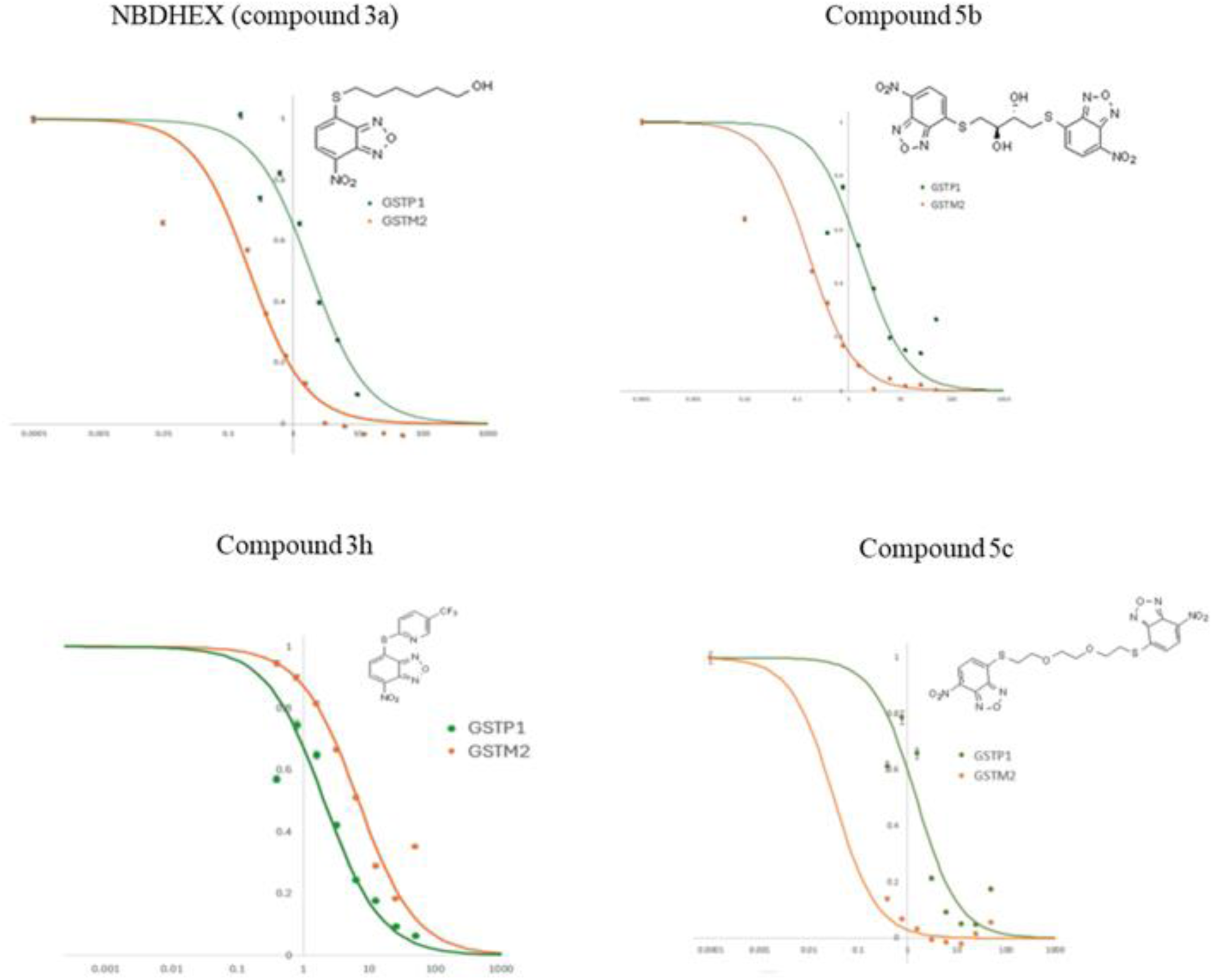
Dose-response curves illustrating the inhibitory effects of the respective compounds on GSTP1 and GSTM2 activity.

**Figure S4:**
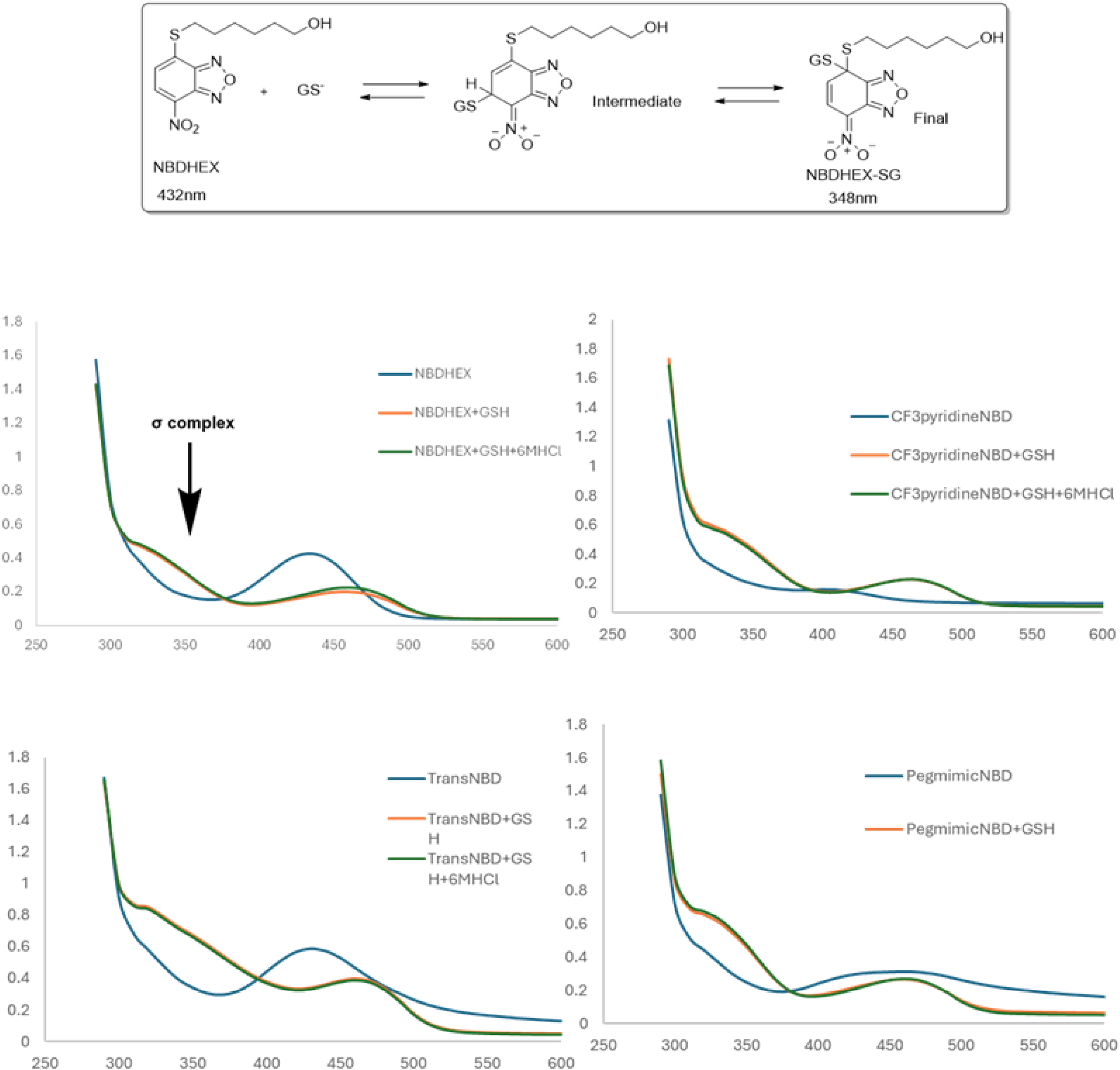
UV-Vis absorption spectra of NBD-based compounds in the presence and absence of glutathione (GSH) and (6M HCl). Spectral analysis of four NBD analogues was performed to evaluate their interaction with GSH and the potential influence of 6MHCl. NBDHEX (**3a),3h, 5b**and **5c**. For each compound, absorption spectra were recorded under three conditions: compound alone (blue line), compound with GSH (orange line), and compound with GSH plus 6MHCl (green line). The addition of GSH resulted in subtle changes in absorption profiles, suggesting interaction and potential conjugation reactions with GSH. The consistency of spectral shifts in the presence of 6MHCl indicates the stability of the resulted complexes. All spectra were recorded from 250 to 600 nm at room temperature.

**Supplementary Figure S5:**
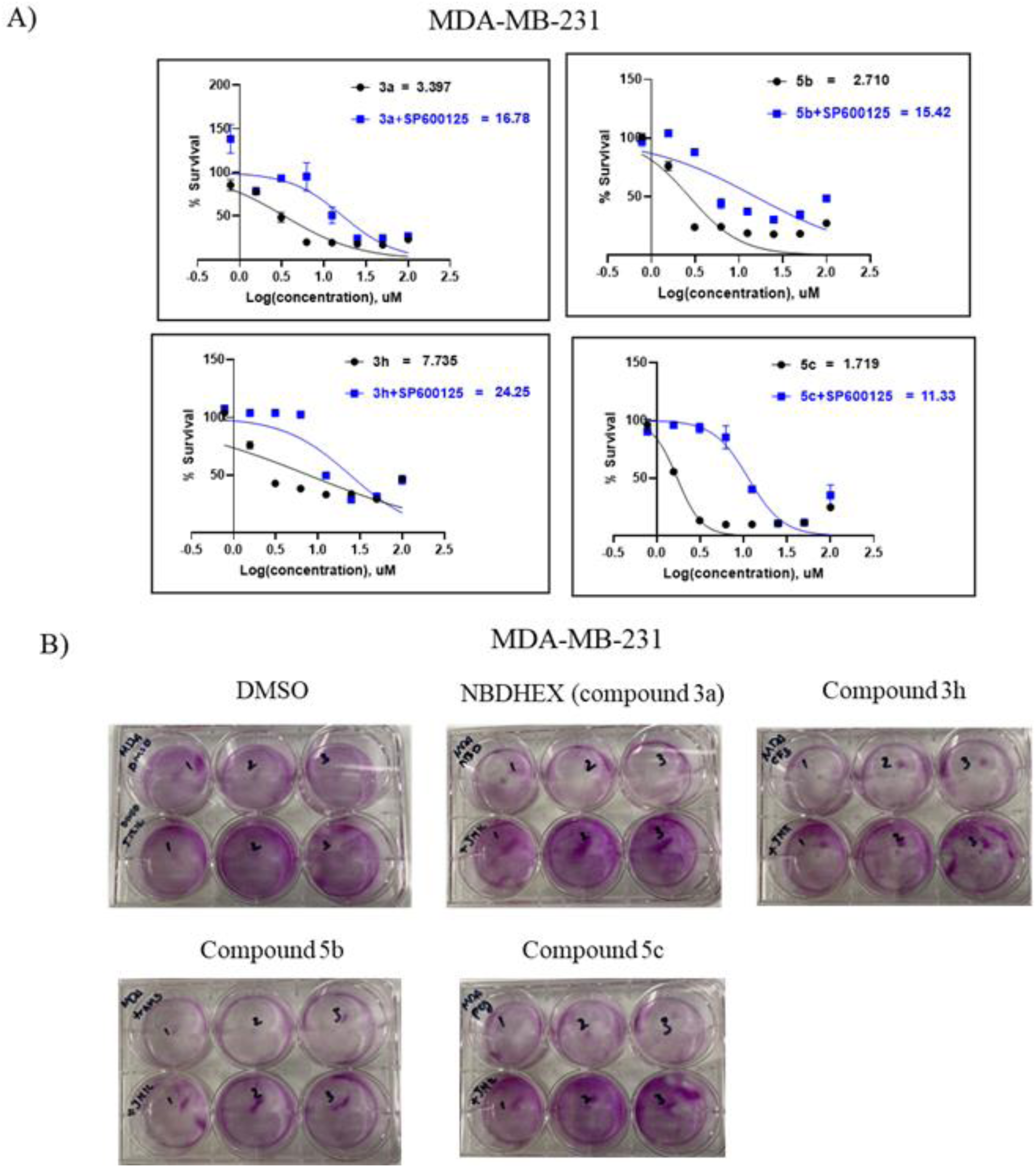

**Figure S6:**
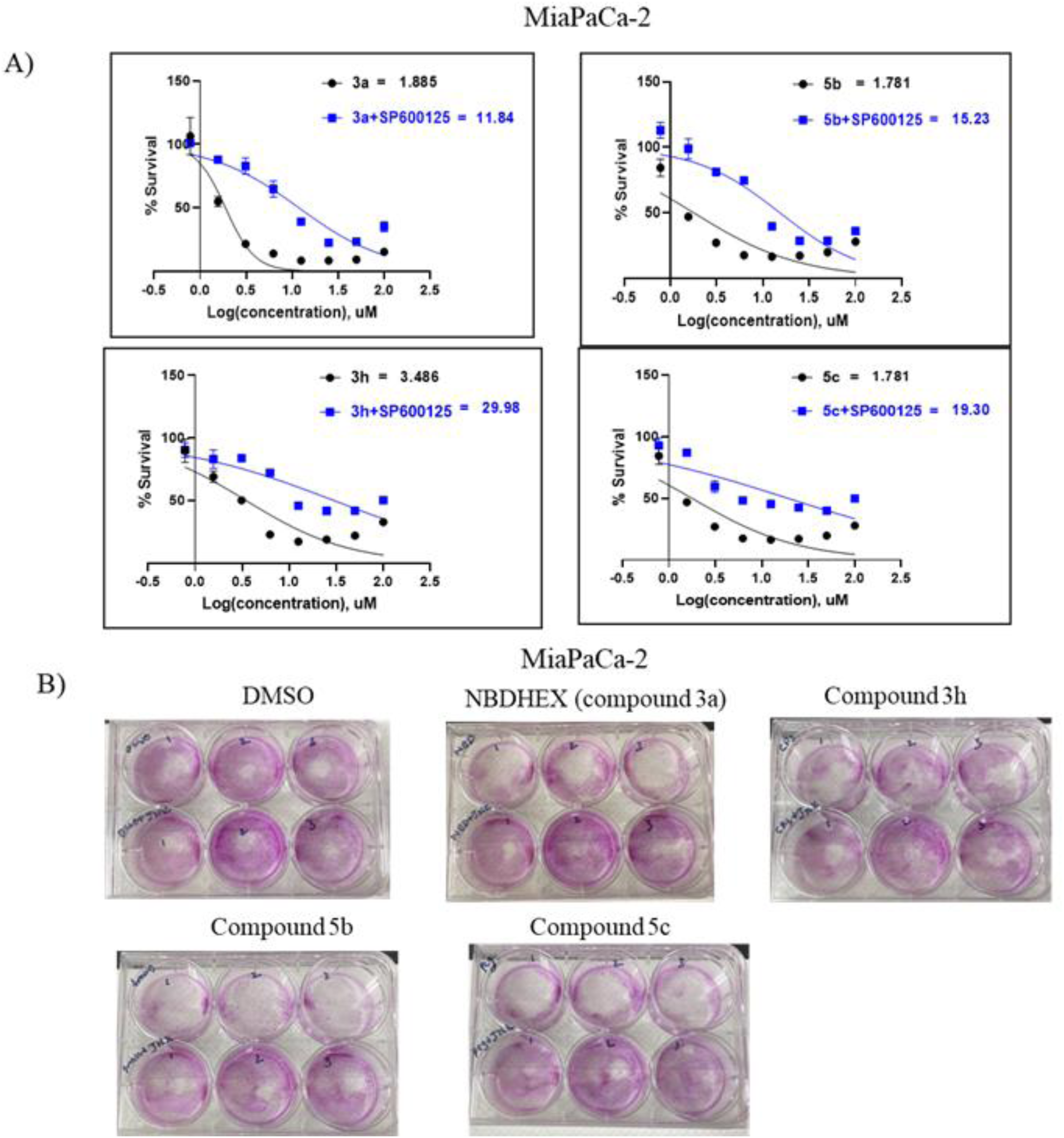
JNK-dependent cytotoxicity of NBDHEX and analogues in MiaPaCa-2 cells**. (A)** Dose-response curves showing cell viability following treatment with NBDHEX (3a) and analogues (3h, 5b, 5c) with or without the JNK inhibitor SP600125. Co-treatment with SP600125 markedly increased IC50 values, indicating that compound-induced cytotoxicity is largely mediated through JNK activation. **(B)** Representative SRB assay images showing reduced colony formation in cells treated with NBDHEX and its analogues compared to DMSO control. Compounds 5b and 5c showed stronger growth inhibition than the parent compound, confirming enhanced cytotoxic potential.

**Figure S7.**
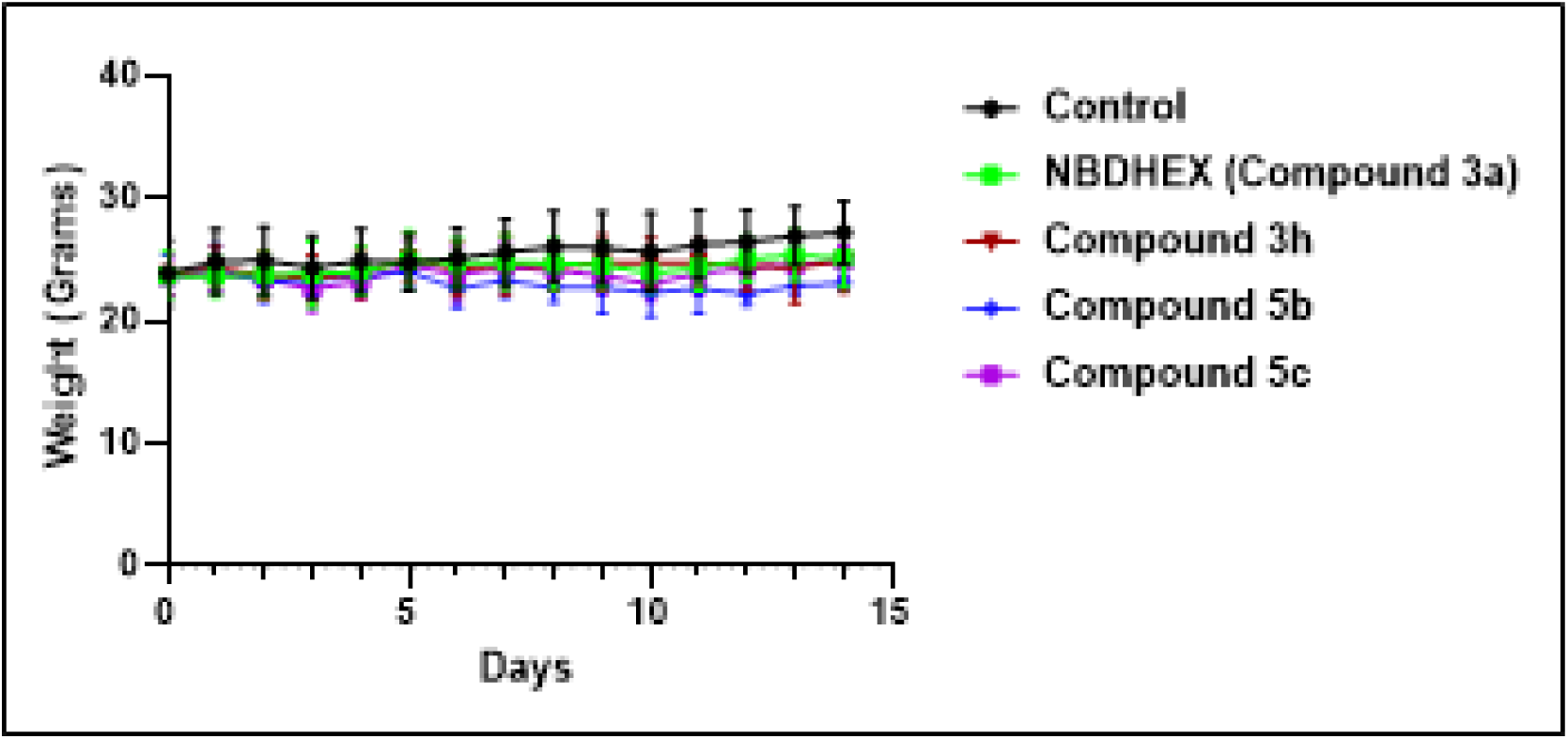
Maximum tolerated dose of compounds **3a,3h,5b** and **5c** administered intraperitoneally and compared with the control. Body weights (g) ± SEM versus day (15 days) after treatment with respective compounds via intraperitoneal injection at 5 mg/kg once daily in healthy CD-1 mice (n=6). Study was carried out under IACUC protocol 2211-40546A.

## APPENDIX

### Spectral characterization

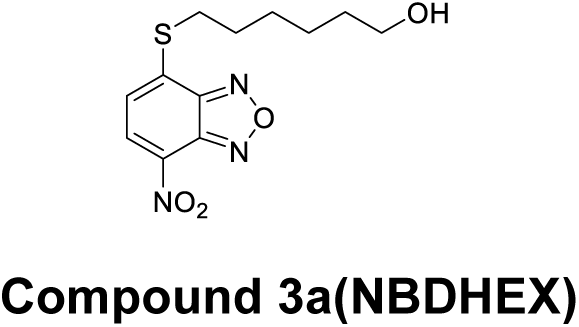

**^1^H NMR (400MHz, Dimethyl Sulfoxide-d_6_):** δ 8.57 (d, J=7.96 Hz, 1H), 7.51 (d, J=8 Hz, 1H), 4.35(t,1H), 3.41-3.34(m, 4H), 1.80-1.73 (m,2H), 1.51-1.31(m,6H)

**^13^ C NMR (100MHz, Dimethyl Sulfoxide-d_6_):** δ 149.68, 143.17, 140.50, 132.81,132.60, 122.67, 61.04, 32.80, 31.11, 28.58, 27.97, 25.48

**HRMS (ESI) m/z:** Calculated for C12H15N3O4S [M]+nNa : 320.068, found at 320.067

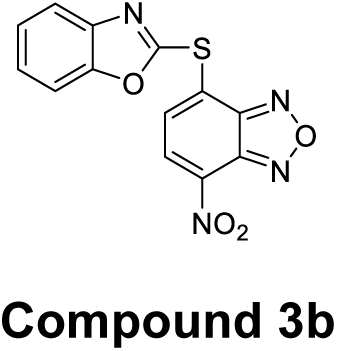

**^1^H NMR (400MHz, Dimethyl Sulfoxide-d_6_):** δ 8.70 (d, J=8Hz, 1H), 8.18 (d, J=7.6Hz, 1H), 7.79-7.74 (m, 2H), 7.48-7.45 (m, 2H)

**^13^ C NMR (100MHz, Dimethyl Sulfoxide-d_6_):** δ 157.26, 151.97, 150.11, 143.51,141.4, 136.7, 133.26, 132.30, 128.09, 126.68, 125.82, 120.13, 111.44

**HRMS (ESI) m/z:** Calculated for C13H6N4O4S [M]+: 315.018, found at 315.024

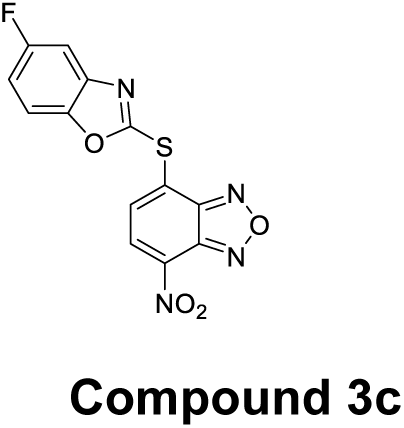

**^1^H NMR (400MHz, Dimethyl Sulfoxide-d_6_):** δ 8.71 (d, J=8Hz, 1H), 8.22 (d, J=8Hz, 1H), 7.80-7.77 (dd, J=8Hz & 4Hz, 1H), 7.69-7.66 (dd, J=4Hz & 4Hz, 1H), 7.69-7.66 (m, 1H)

**^13^ C NMR (100MHz, Dimethyl Sulfoxide-d_6_):** δ 161.32, 159.65, 158.94, 150.22, 148.52, 143.51, 142.27 (d, J=13.61Hz), 134.10, 132.23, 127.37, 114.12 (d, J=26.19Hz), 112.35 (d, J=10.17Hz), 106.59 (d, J=26.1Hz)

**HRMS (ESI) m/z:** Calculated for C13H5FN4O4S [M]+: 333.009, found at 333.021

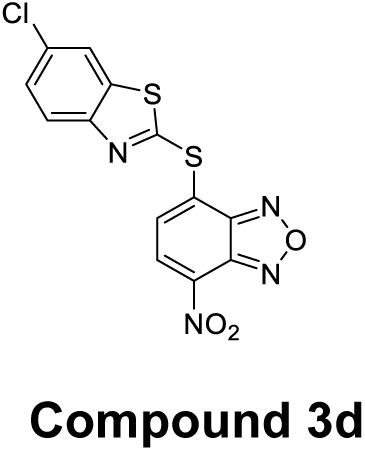

**^1^H NMR (400MHz, Dimethyl Sulfoxide-d_6_):** δ 8.65 (d, J=8Hz, 1H), 8.32 (d, J=2.11Hz, 1H), 8.01 (d, J=2.21Hz, 1H), 8.00 (d, J=1.94 Hz, 1H), 7.63( dd, 0.55Hz & 1.64Hz, 1H)

**^13^ C NMR (100MHz, Dimethyl Sulfoxide-d_6_):** δ 159.99, 151.88, 150.02, 143.49,138.43, 136.35, 132.37, 132.28, 131.36, 131.13, 128.03, 124.46, 122.56

**HRMS (ESI) m/z:** Calculated for C13H5ClN4O3S2 [M]+: 364.956, found at 364.961

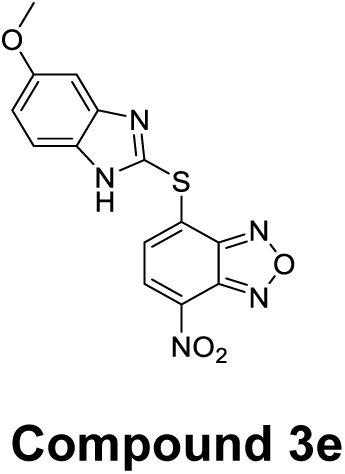

**^1^H NMR (400MHz, Dimethyl Sulfoxide-d_6_):** δ 8.57 (d, J=7.88 Hz, 1H), 7.57 (d, J=8.84 Hz, 1H), 7.39 (d, J=7.92 Hz,1H), 7.11(s, 1H), 6.94 (dd, J=2.4Hz & 6.48Hz 1H), 3.82 (s,3H)

**^13^ C NMR (100MHz, Dimethyl Sulfoxide-d_6_):** δ 157.13, 149.21, 143.35, 138.23, 135.23, 134.59, 132.81, 126.99, 113.83, 56.04

**HRMS (ESI) m/z:** Calculated for C14H9N5O4S [M]+nH : 344.045, found at 344.057

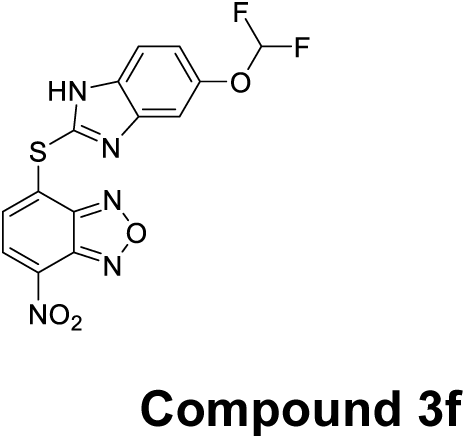

**^1^H NMR (400MHz, Dimethyl Sulfoxide-d_6_):** δ 8.59 (d, J=8Hz, 1H), 7.69 (d, J=8Hz, 1H), 7.55 (d, J=8Hz, 1H), 7.46 (d, J=4Hz, 1H), 7.26 (s,1H), 7.18-7.15 (m, 1H), 7.07 (s,1H)

**^13^ C NMR (100MHz, Dimethyl Sulfoxide-d_6_):** δ 149.45, 147.55, 143.39, 141.78, 135.00, 133.83, 132.72, 130.97, 128.23, 119.78, 117.22, 116.35, 114.66

**HRMS (ESI) m/z:** Calculated for C14H7F2N5O4S [M]+ : 380.026, found at 380.041

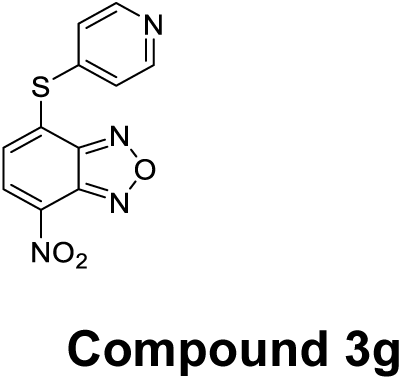

**^1^H NMR (400MHz, Dimethyl Sulfoxide-d_6_):** δ 8.67-8.66 (m, 2H), 8.58 (d, J=8Hz, 1H), 7.62-7.60 (m, 2H), 7.53 (d, J=8Hz,1H),

**^13^ C NMR (100MHz, Dimethyl Sulfoxide-d_6_):** δ 151.25, 150.05, 143.57, 140.55, 135.37, 133.95, 132.62, 130.14, 126.72

**HRMS (ESI) m/z:** Calculated for C11H6N4O3S [M]+ : 275.023, found at 275.023

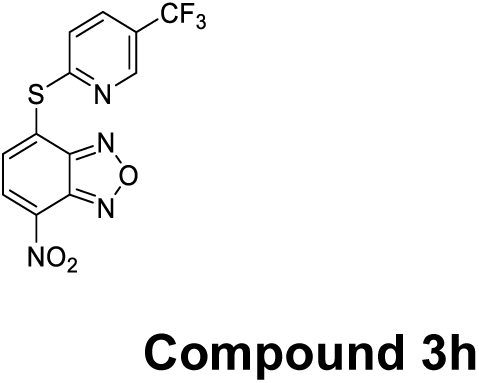

**^1^H NMR (400MHz, Dimethyl Sulfoxide-d_6_):** δ 8.87 (s, 1H), 8.67 (d, J=8Hz, 1H), 8.24(d, J=4Hz, 1H), 8.06 (d, J=8 Hz, 1H), 7.85 (d, J=8Hz, 1H)

**^13^ C NMR (100MHz, Dimethyl Sulfoxide-d_6_):** δ 159.77, 150.76, 147.33 (q, J=4.14Hz), 143.6, 136.55, 135.76 (q, J=3.23Hz), 134.29, 132.37, 130.51, 125.24, 124.30 (q, J=32.59Hz), 122.57

**HRMS (ESI) m/z:** Calculated for C12H5F3N4O3S [M]+ : 343.011, found at 343.022

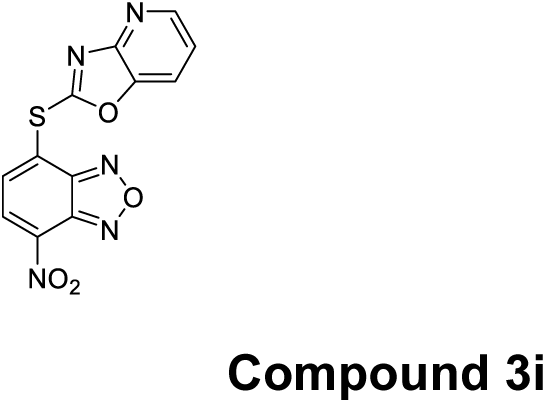

**^1^H NMR (400MHz, Dimethyl Sulfoxide-d_6_):** δ 8.76 (d, J=8Hz, 1H), 8.55 (d, J=8Hz, 1H), 8.37 (d, J=8Hz, 1H), 8.21-8.18 (m, 1H) 7.51-7.48 (m, 1H)

**^13^ C NMR (100MHz, Dimethyl Sulfoxide-d_6_):** δ 162.17, 154.98, 150.41, 147.33,144.48, 143.52, 137.23, 135.19, 132.21, 126.34, 121.68, 119.64

**HRMS (ESI) m/z**: Calculated for C12H5N5O4S [M]+ : 316.014, found at 316.019

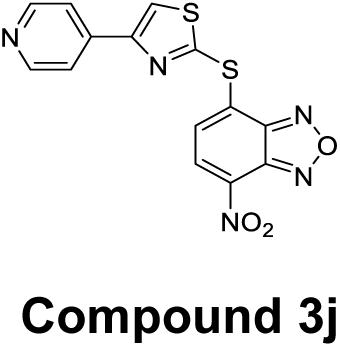

**^1^H NMR (400MHz, Dimethyl Sulfoxide-d_6_):** δ 8.89 (s, 1H), 8.70 (s, 2H), 8.59 (d, J=4Hz, 1H), 7.98 (s, 2H), 7.62 (d, J=8 Hz, 1H)

**^13^ C NMR (100MHz, Dimethyl Sulfoxide-d_6_):** δ 154.57,154.13,150.57,149.3,143.39,140.6, 135.20, 134.53, 132.68, 128.09, 125.80, 120.93, 56.5, 19.02

**HRMS (ESI) m/z:** Calculated for C14H7N5O3S2 [M]+ : 358.006, found at 358.017

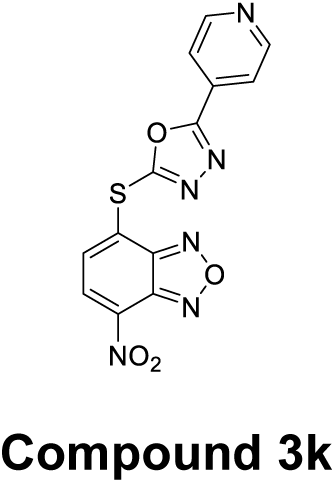

**^1^H NMR (400MHz, Dimethyl Sulfoxide-d_6_):** δ 8.86 (d, J=4.4Hz, 2H), 8.65 (d, J=8Hz, 1H), 8.08 (d, J=7.6Hz, 1H), 7.95 (d, J=4 Hz, 2H)

**^13^ C NMR (100MHz, Dimethyl Sulfoxide-d_6_):** δ 165.94, 159.27, 151.55, 149.74, 143.50, 136.50, 132.27, 132.13, 130.27, 128.40, 120.80

**HRMS (ESI) m/z:** Calculated for C13H6N6O4S [M]+ : 343.024, found at 343.031

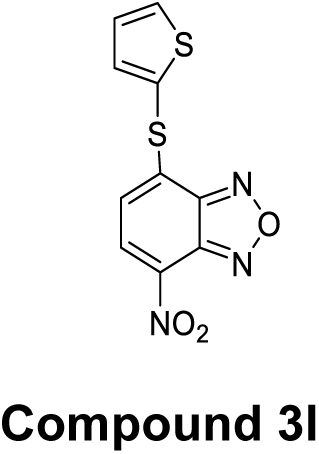

**^1^H NMR (400MHz, Dimethyl Sulfoxide-d_6_):** δ 8.56 (d, J=8Hz, 1H), 8.15-8.14 (dd, J=1.24Hz & 4.12Hz, 1H), 7.71-7.70 (dd, J=8 & 1.2Hz, 1H), 7.41-7.39 (dd, J=8 &1.2Hz, 1H), 6.9 (d, J=4Hz,1H)

**^13^ C NMR (100MHz, Dimethyl Sulfoxide-d_6_):** δ 148.62, 143.25, 140.47, 140.06, 136.33,133.78, 132.91, 130.04, 123.92, 121.97

**HRMS (ESI) m/z:** Calculated for C13H6N4O4S [M]+ : 279.985, found at 280.007

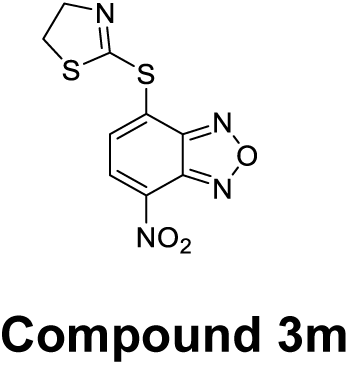

**^1^H NMR (400MHz, Dimethyl Sulfoxide-d_6_):** δ 8.68 (d, J=8Hz, 1H), 8.29 (d, J=8Hz, 1H), 4.24(t, J=8Hz, 2H), 3.54 (t, 8 Hz, 2H)

**^13^ C NMR (100MHz, Dimethyl Sulfoxide-d_6_):** δ 159.75, 150.58, 143.3, 136.71, 134.55, 132.14, 129.06, 65.14, 35.99

**HRMS (ESI) m/z:** Calculated for C9H6N4O3S2 [M]+ : 282.995, found at 283.020

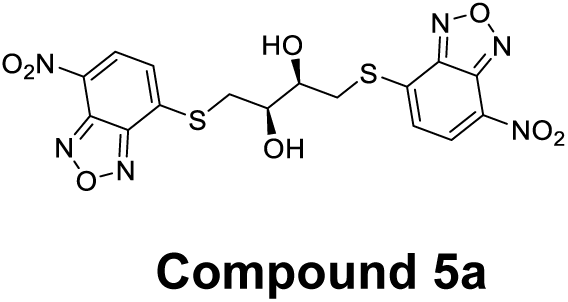

**^1^H NMR (400MHz, Dimethyl Sulfoxide-d_6_):** δ 8.59 (d, J=7.84 Hz, 2H), 7.57 (d, J=8.04 Hz, 2H), 5.69 (s, 2H), 3.96 (s, 2H), 3.62-3.58 (m, 2H), 3.51-3.45 (m, 2H)

**^13^ C NMR (100MHz, Dimethyl Sulfoxide-d_6_):** δ 149.68, 143.08, 140.80, 132.78, 132.59, 122.80, 71.14, 34.85

**HRMS (ESI) m/z:** Calculated for C14H9N5O4S [M]+nH : 481.023, found at 481.038

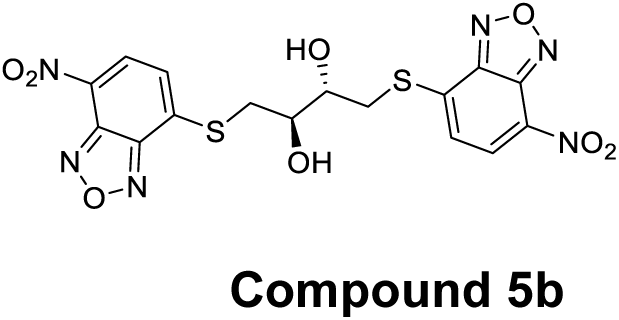

**^1^H NMR (400MHz, Dimethyl Sulfoxide-d_6_):** δ 8.61 (d, J=8 Hz, 2H), 7.56 (d, J=8.08 Hz, 2H), 5.87 (d, J=4.34 Hz, 2H), 3.89 (br, s, 2H), 3.78-3.75 (m, 2H), 3.48-3.43 (m, 2H)

**^13^ C NMR (100MHz, Dimethyl Sulfoxide-d_6_):** δ 149.71, 143.08, 141.11, 132.87, 132.54, 122.71, 72.02, 36.33

**HRMS (ESI) m/z:** Calculated for C14H9N5O4S [M]+nNa: 503.01, found at 503.00

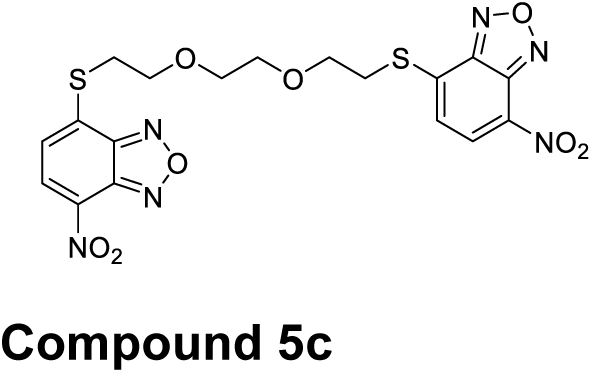

**^1^H NMR (400MHz, Dimethyl Sulfoxide-d_6_):** δ 8.55 (d, J=8 Hz, 2H), 7.55 (d, J=8.08 Hz, 2H), 3.82 (t, J= 8 Hz, 4H), 3.62 (s, 4H), 3.56 (t, J=11Hz, 4H)

**^13^ C NMR (100MHz, Dimethyl Sulfoxide-d_6_):** δ 149.56, 142.99, 140.26, 132.72, 132.58, 122.95, 70.32, 68.34, 31.64

**HRMS (ESI) m/z:** Calculated for C14H9N5O4S [M]+nH: 509.05, found at 509.07

### NMR Spectra

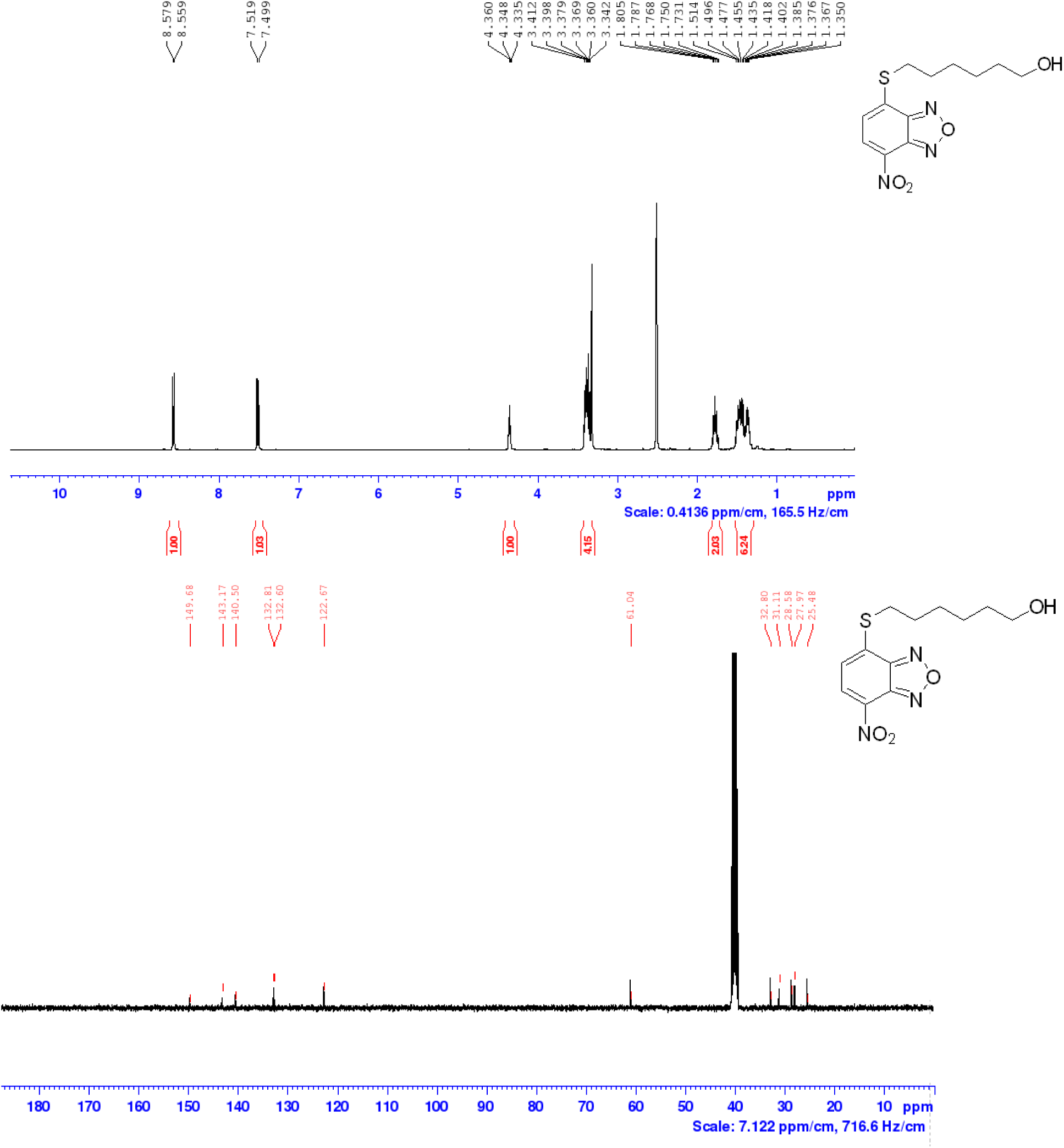

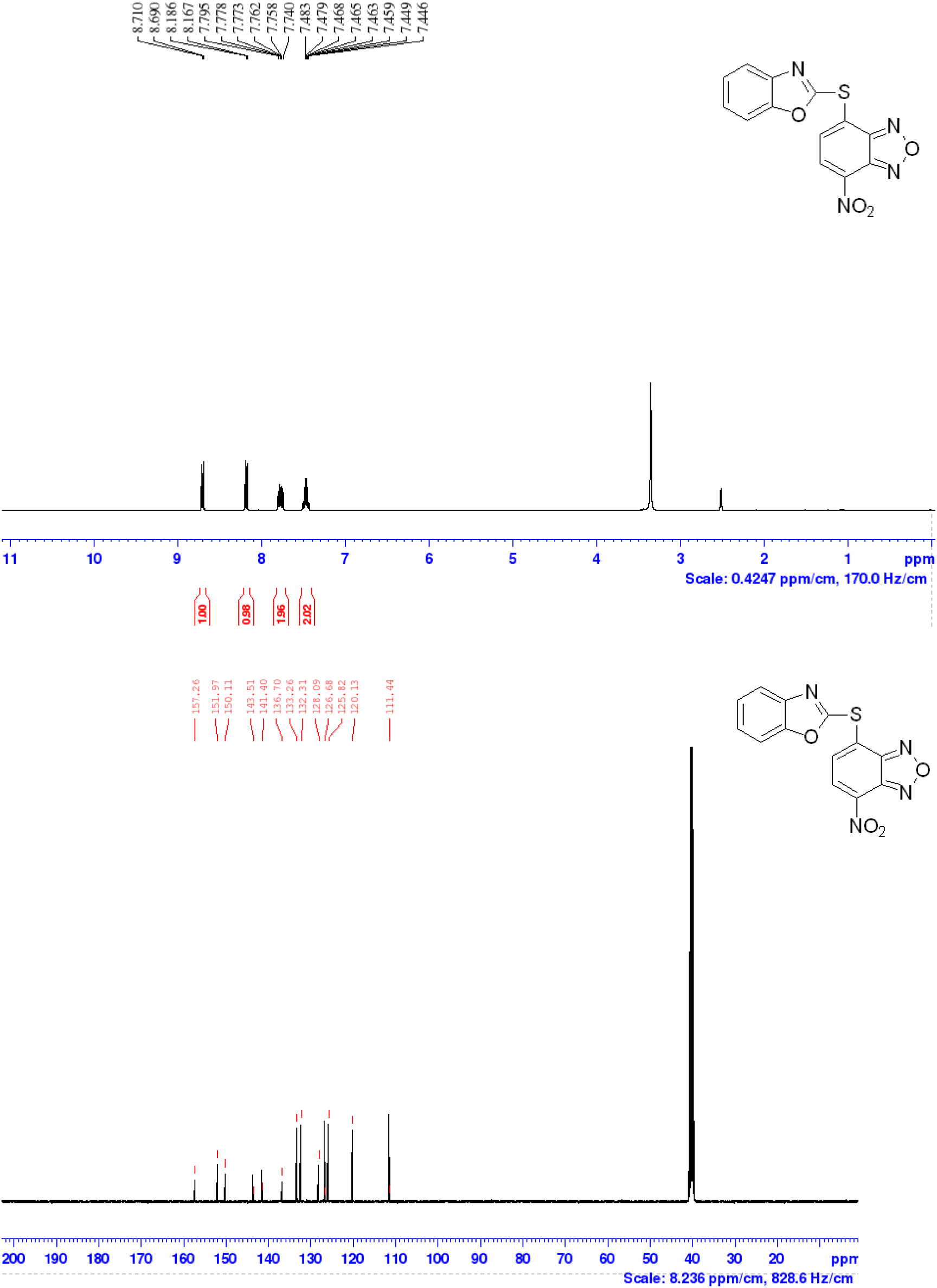

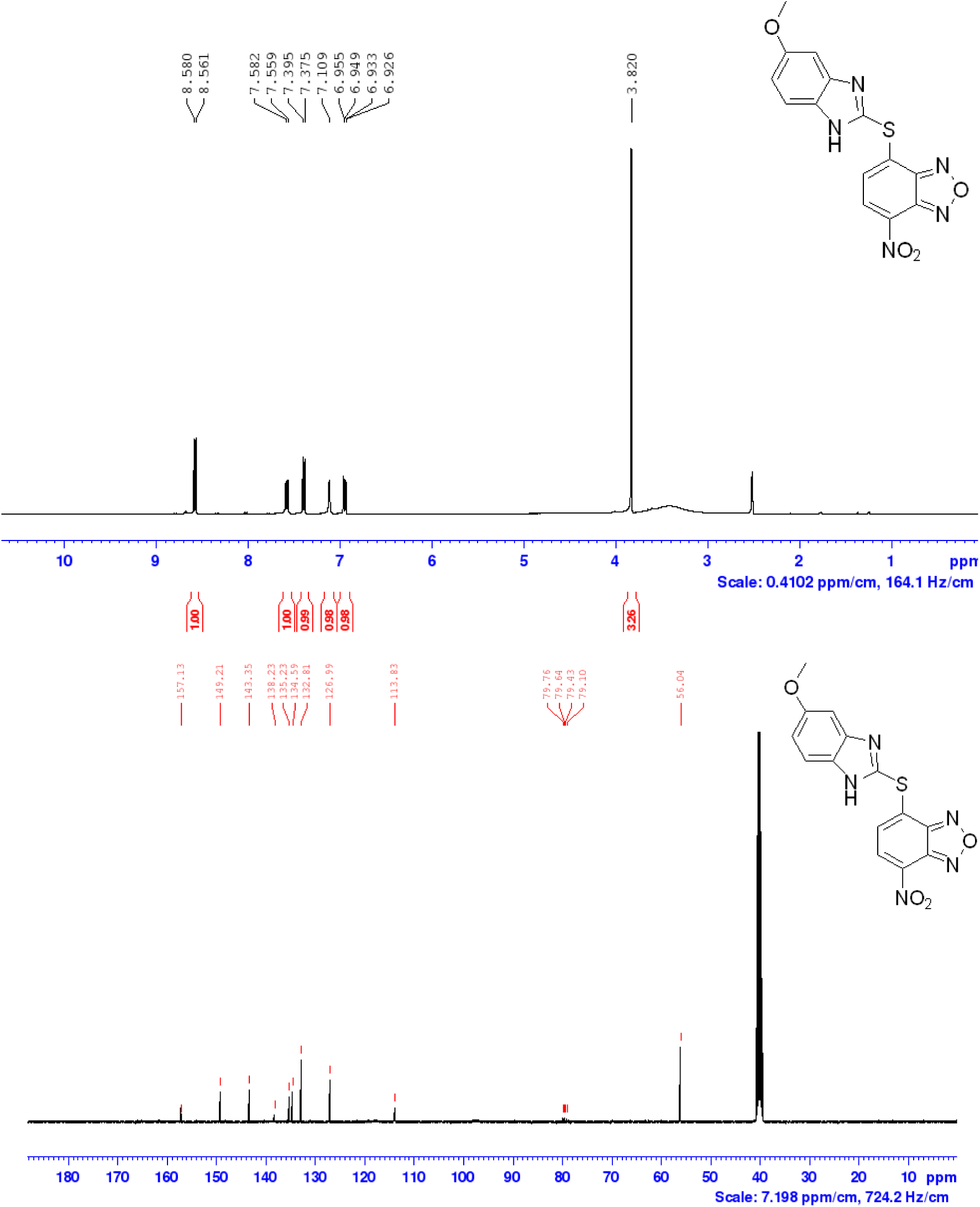

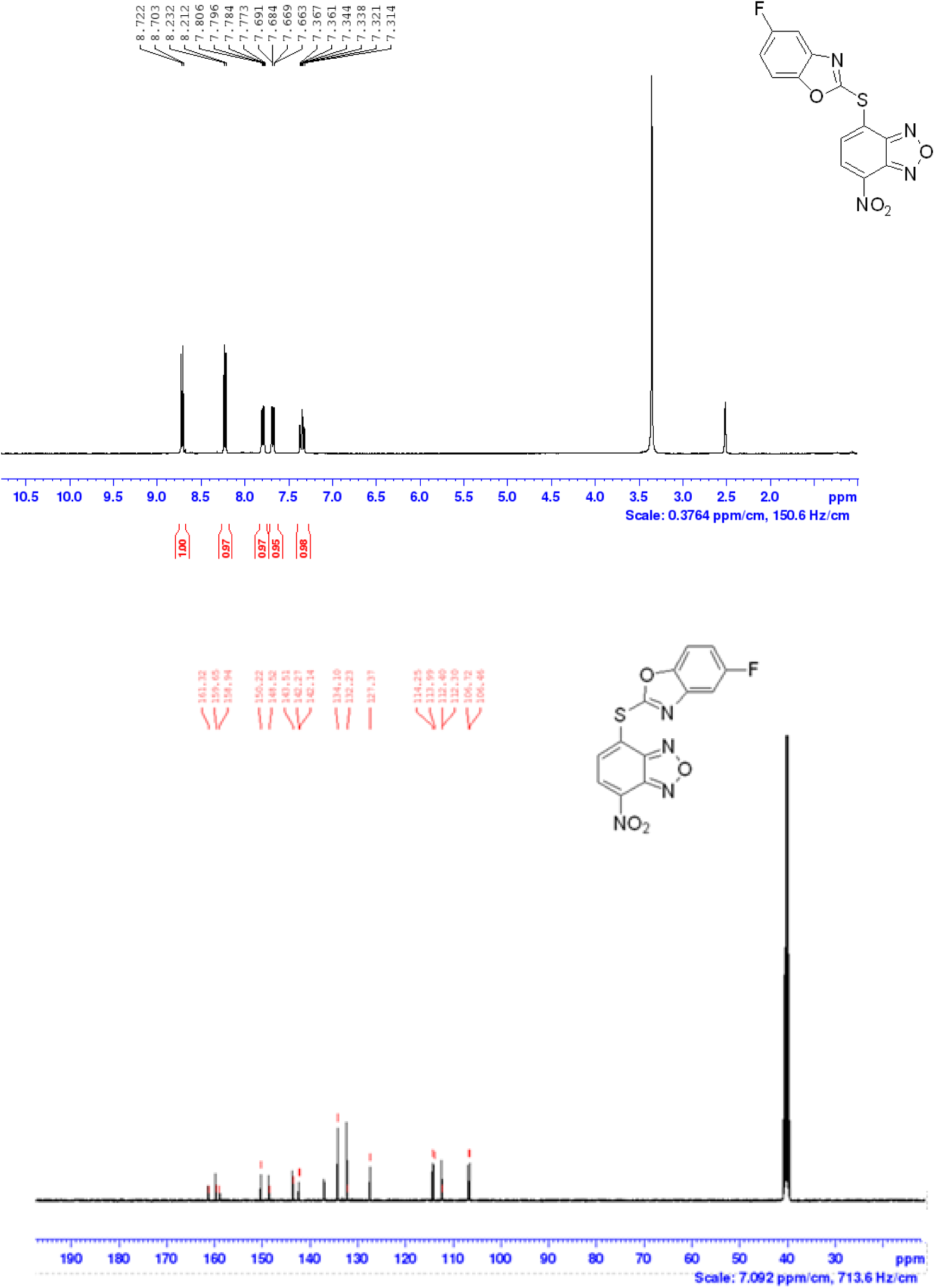

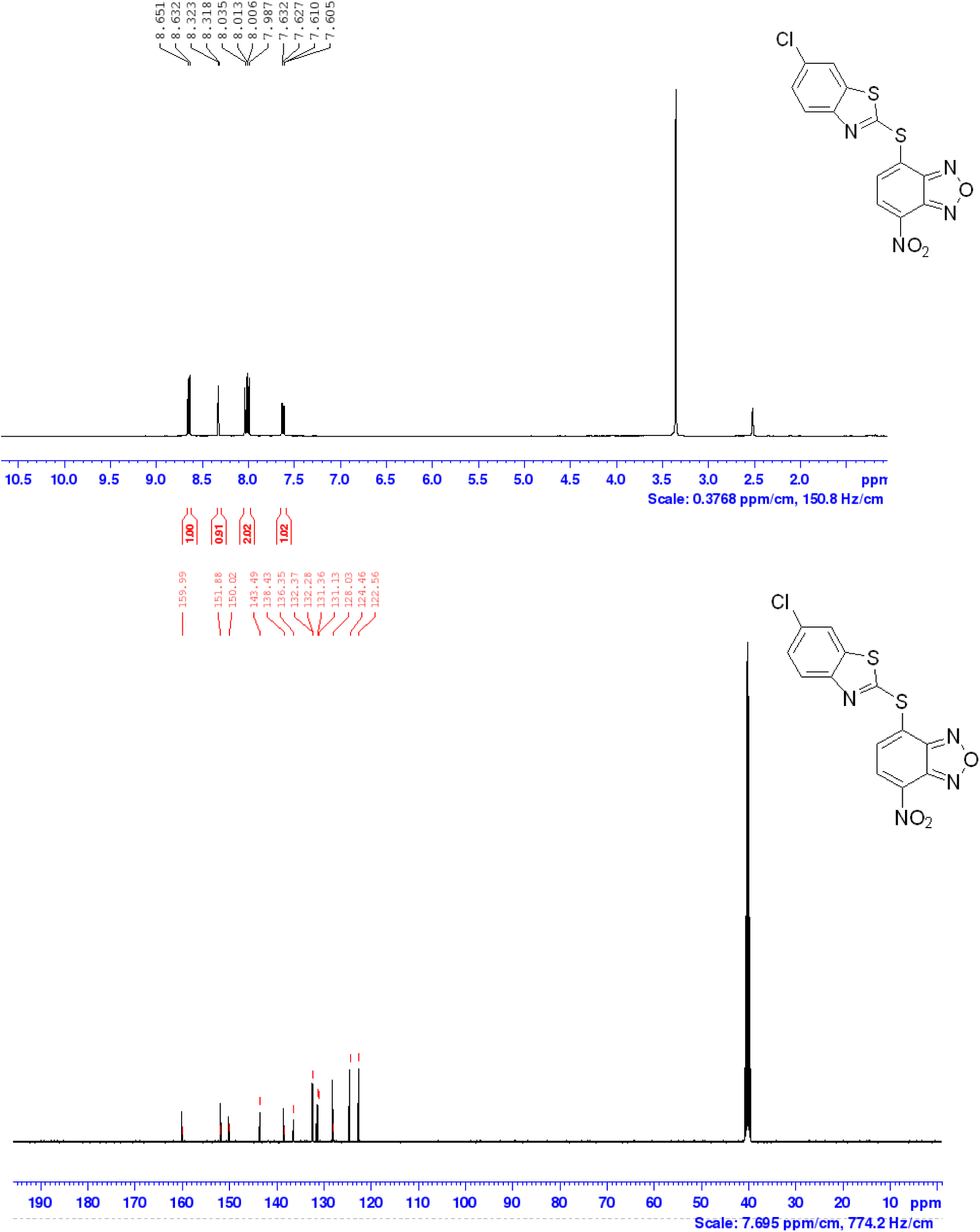

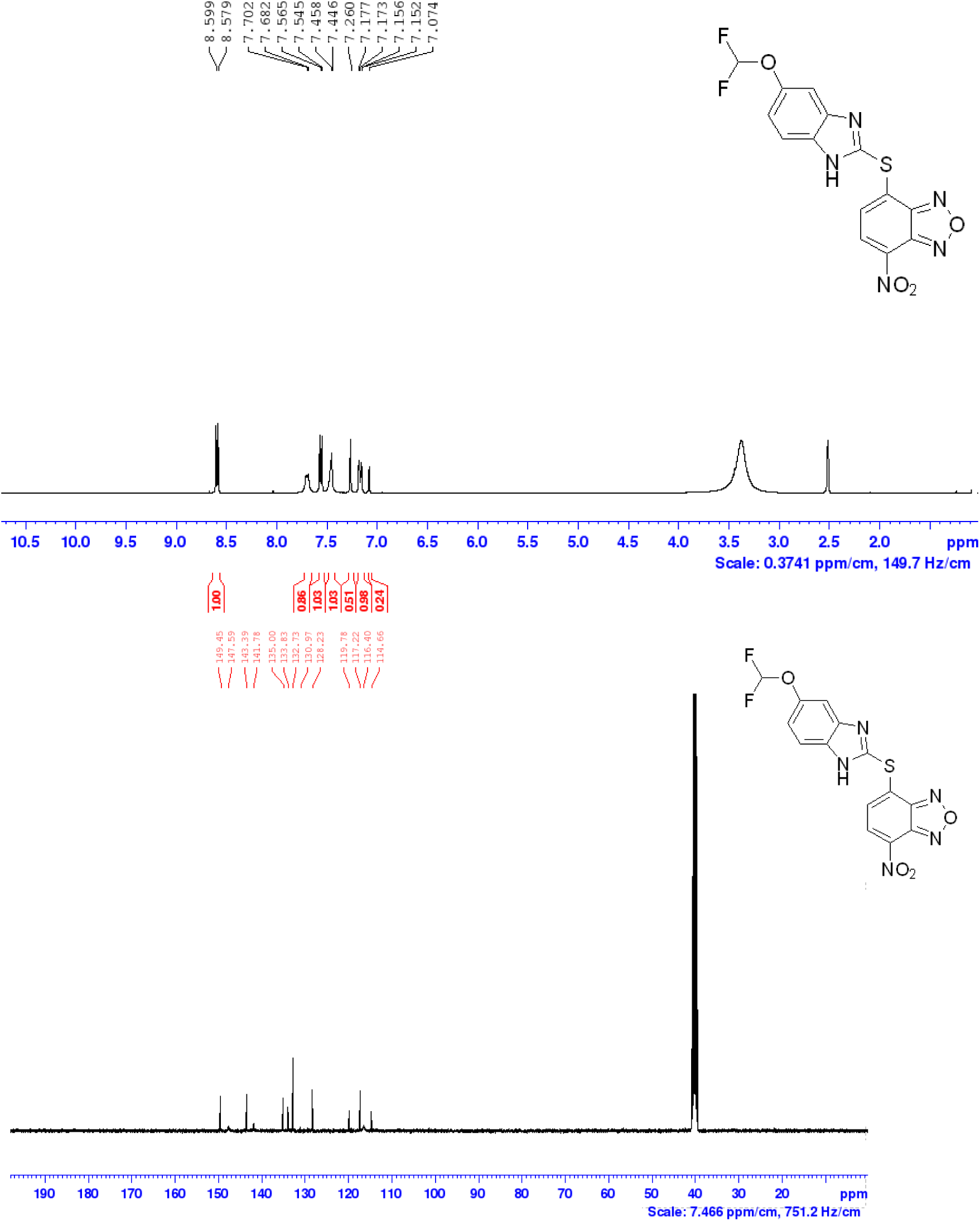

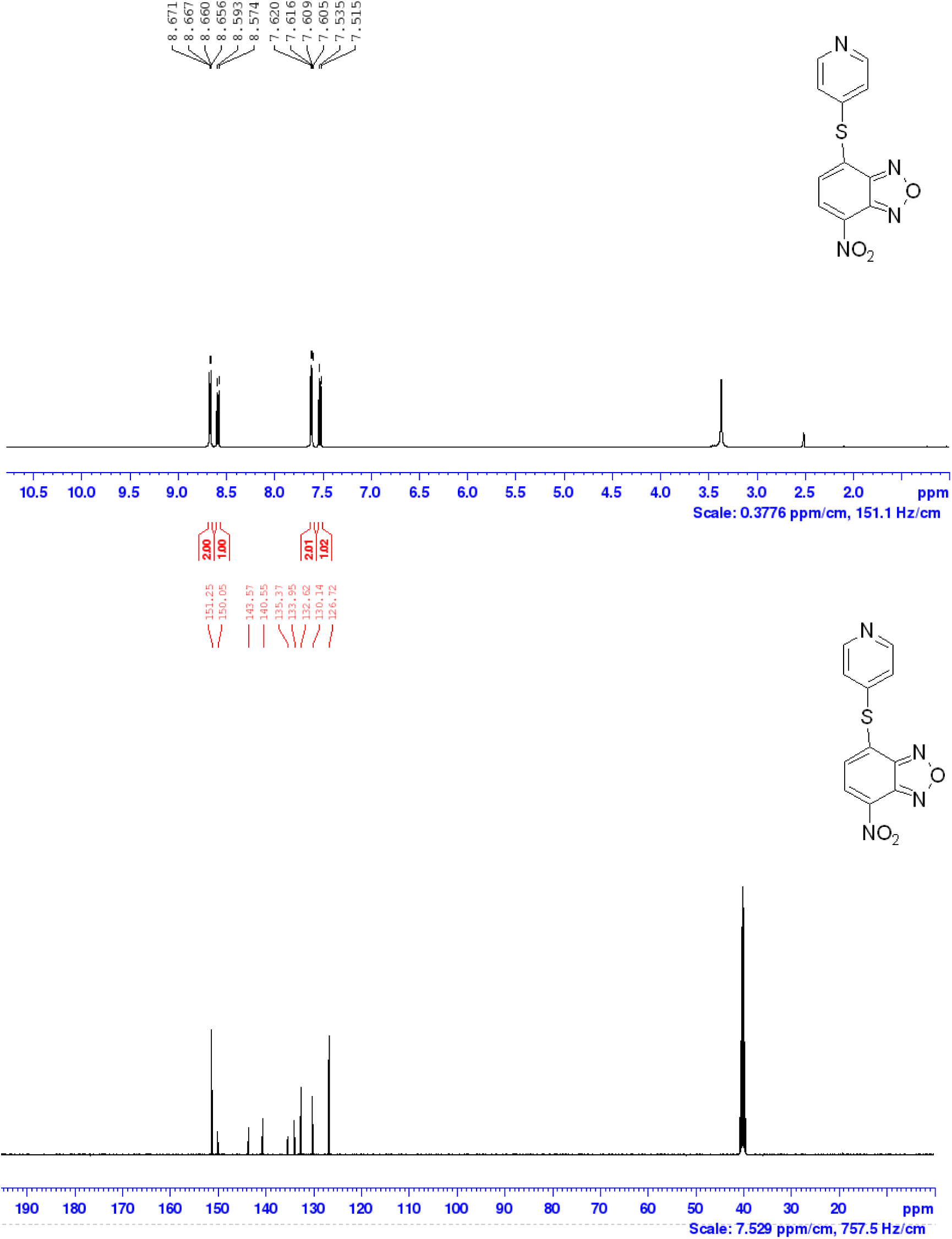

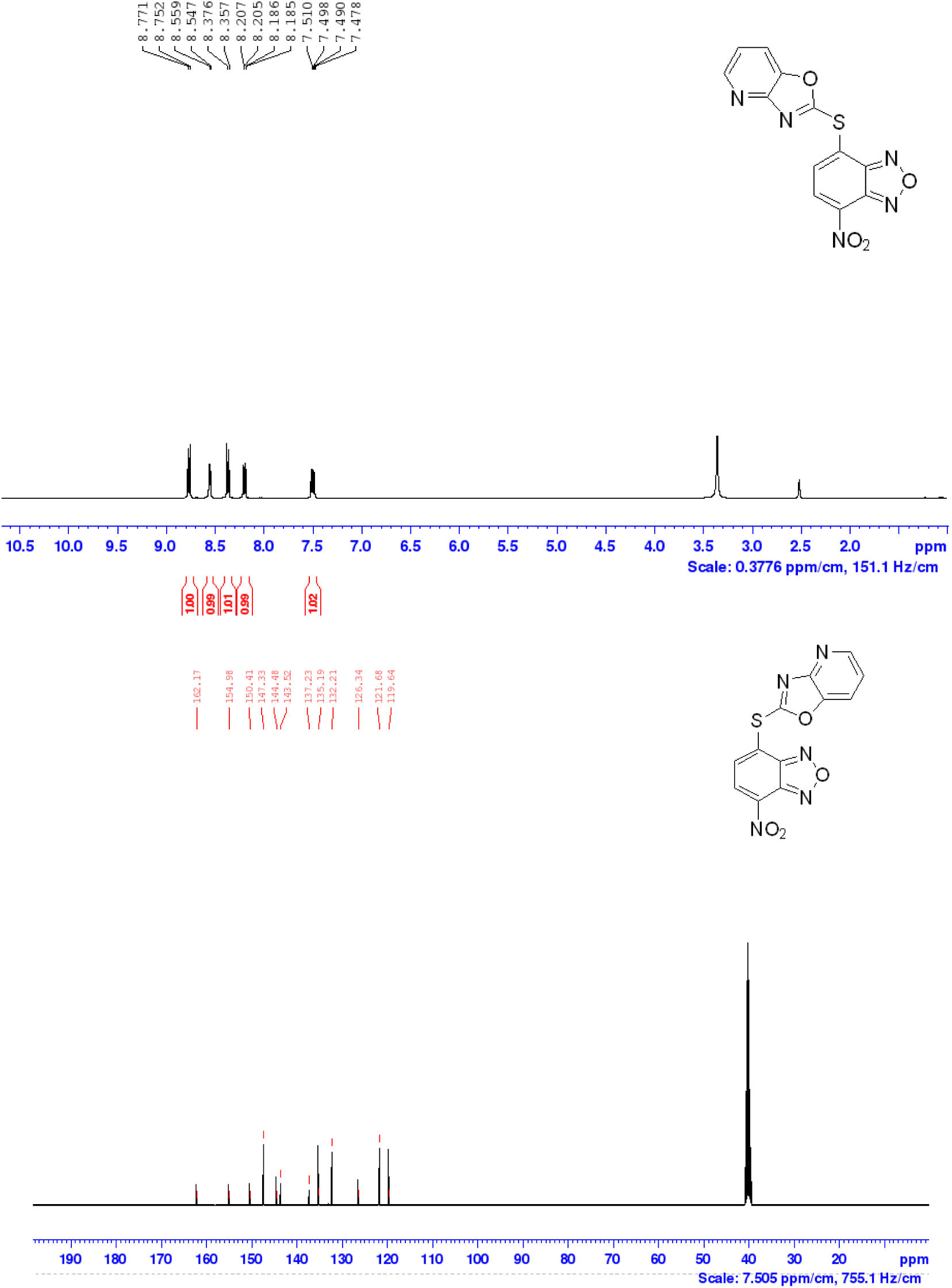

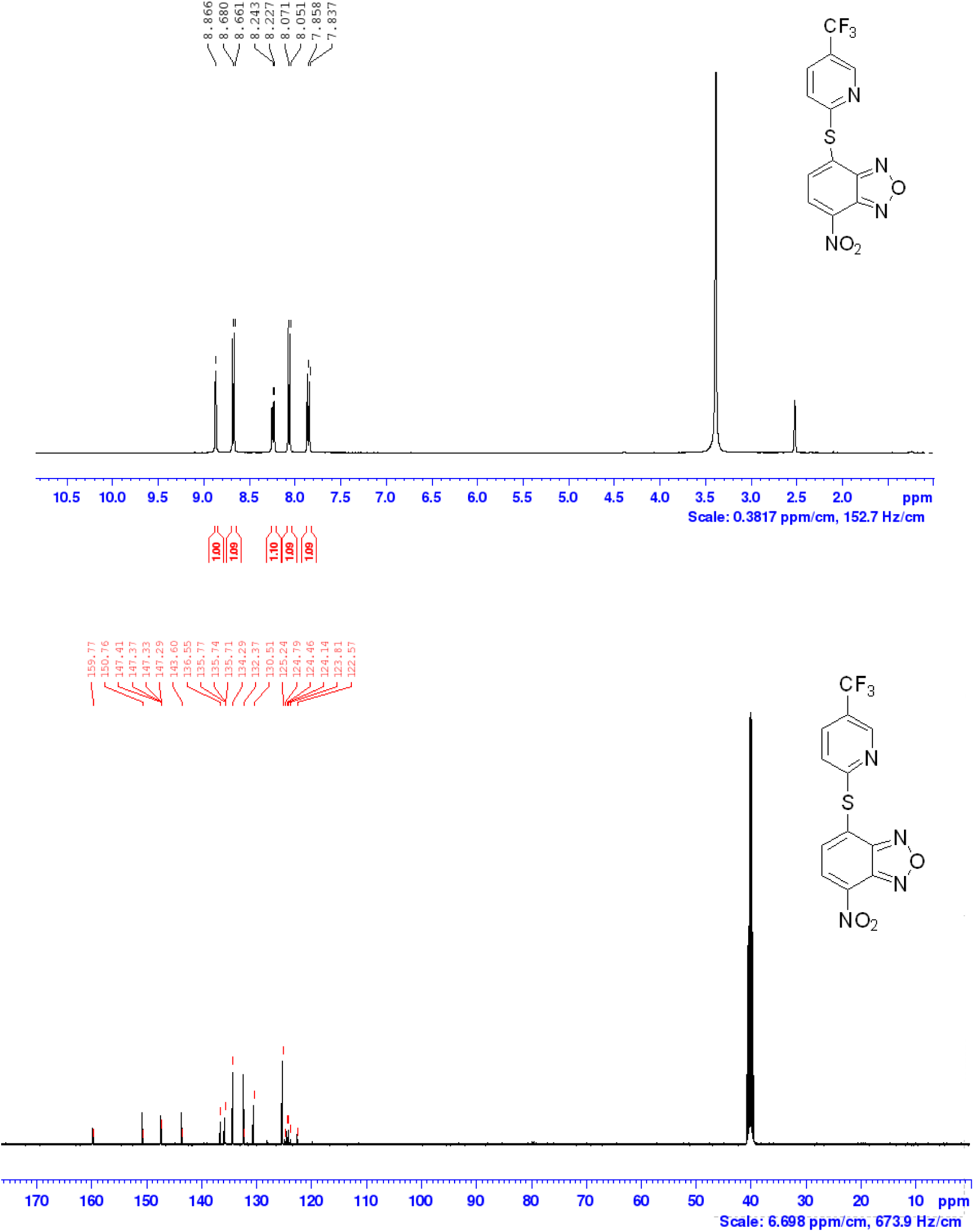

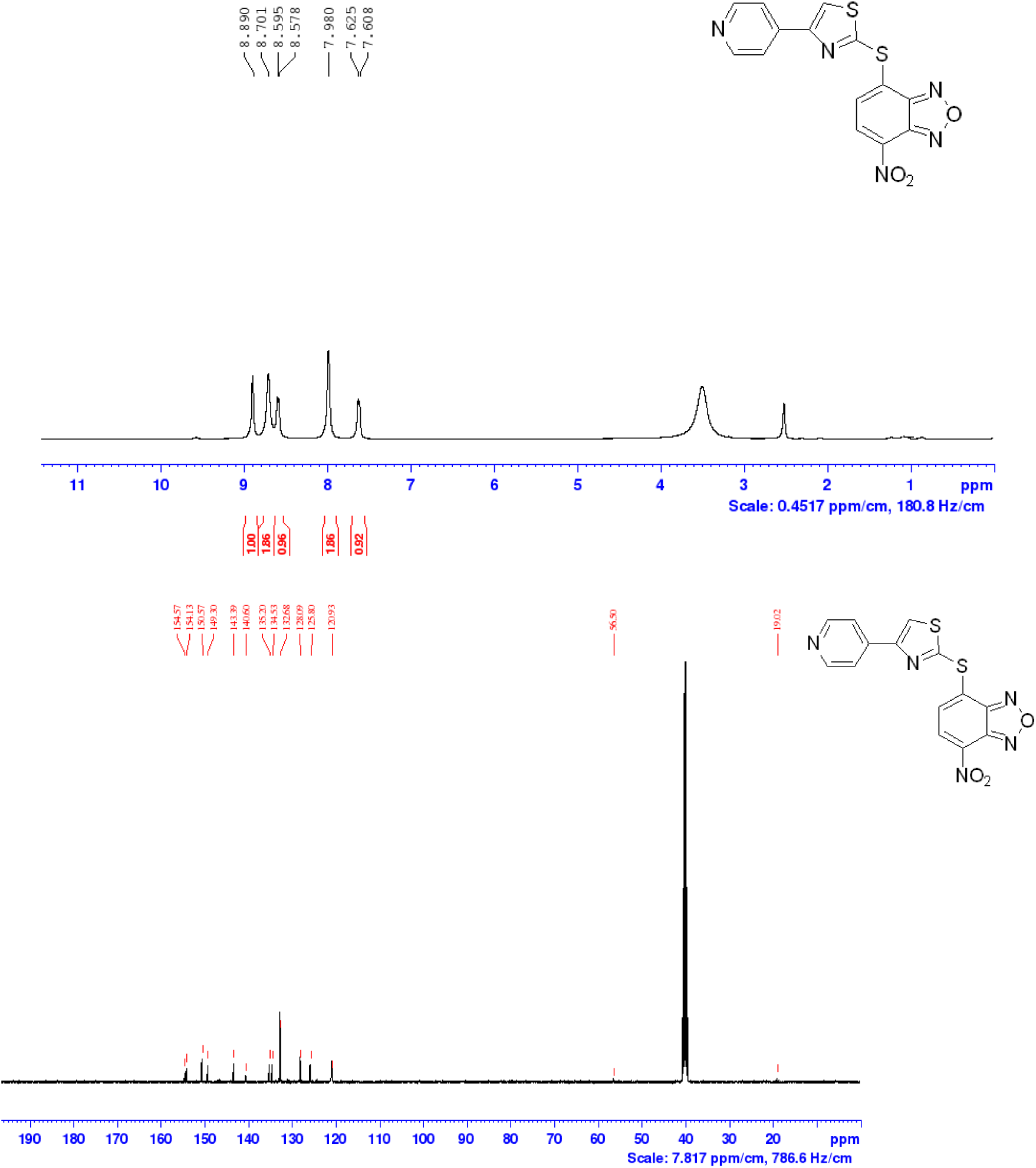

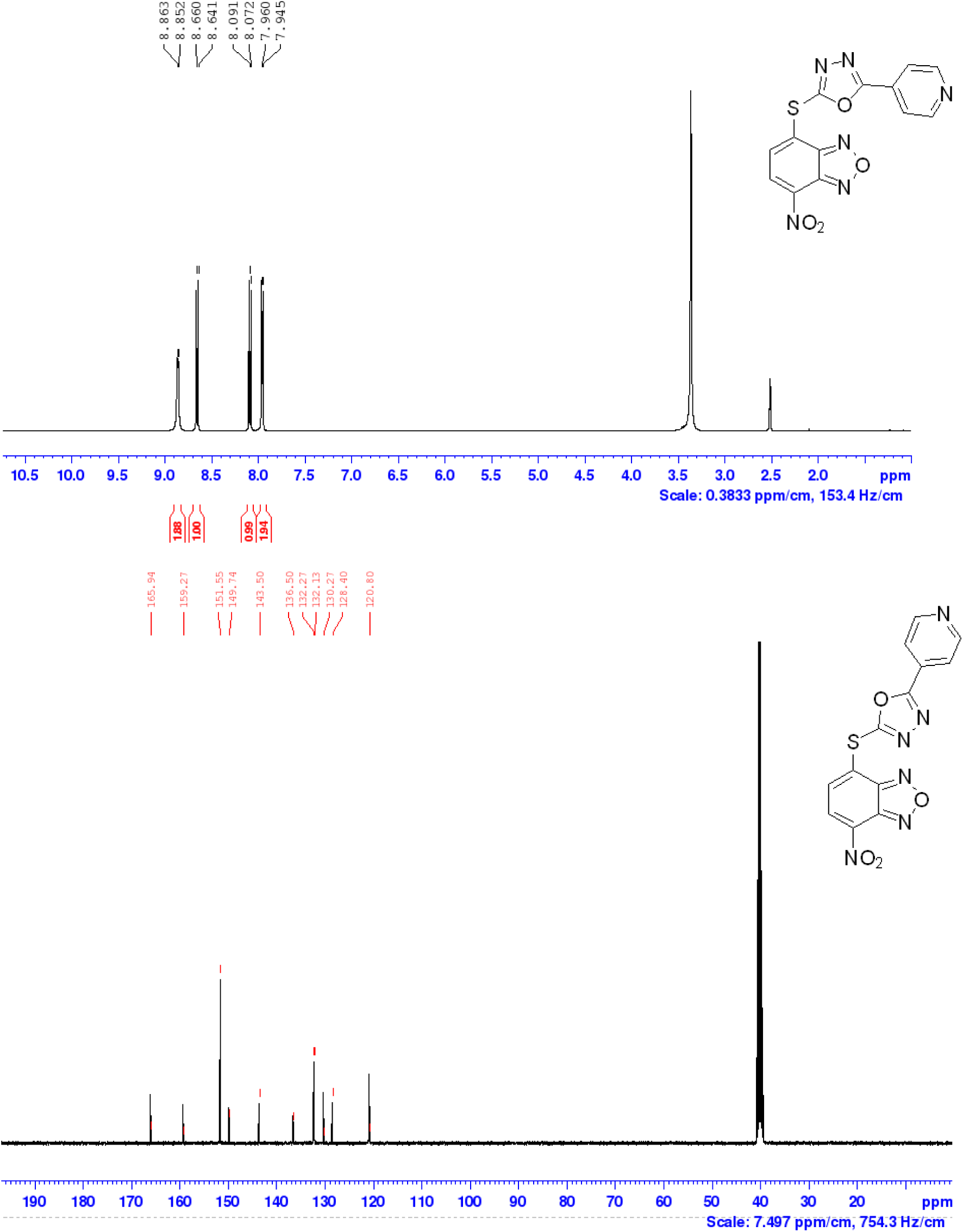

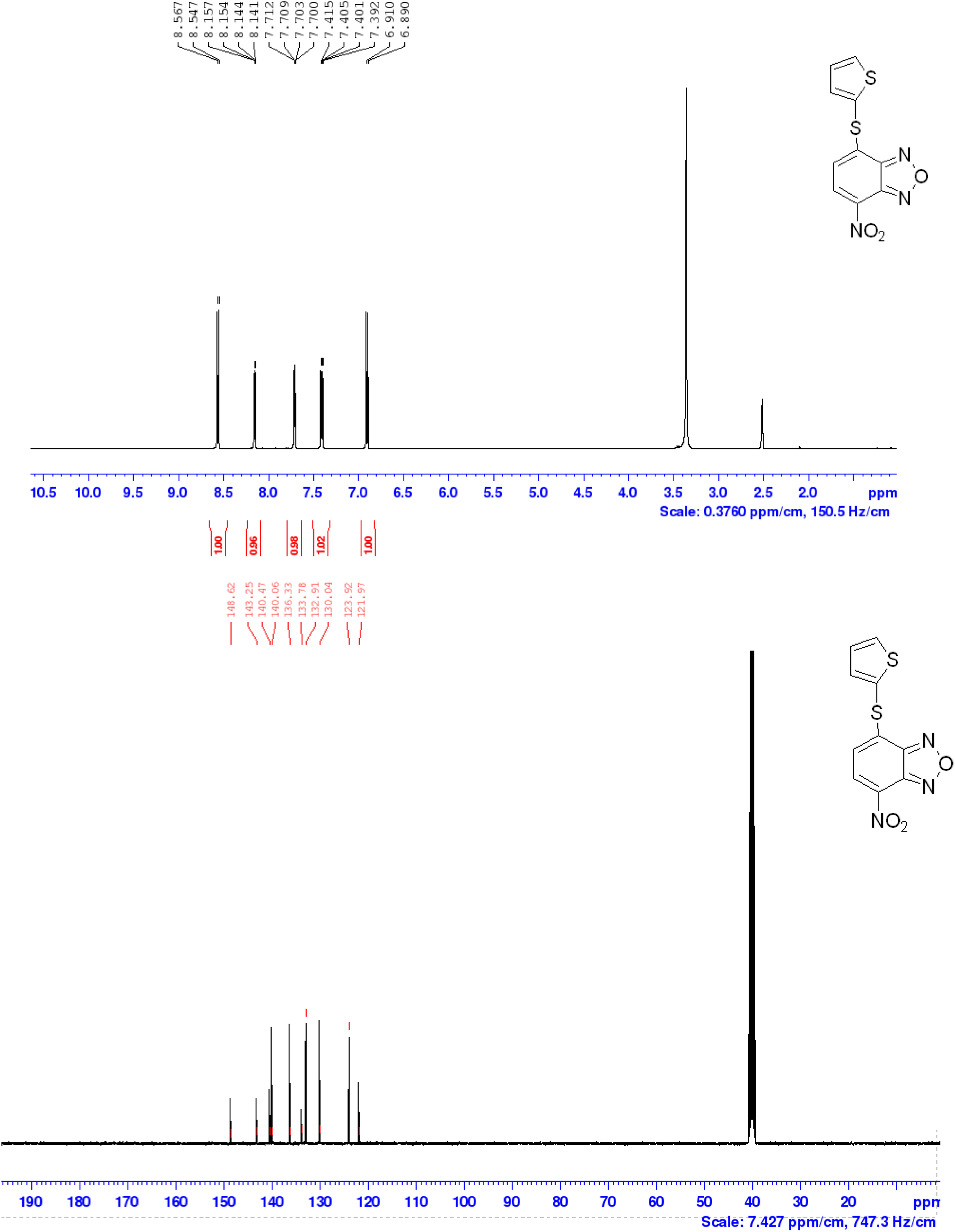

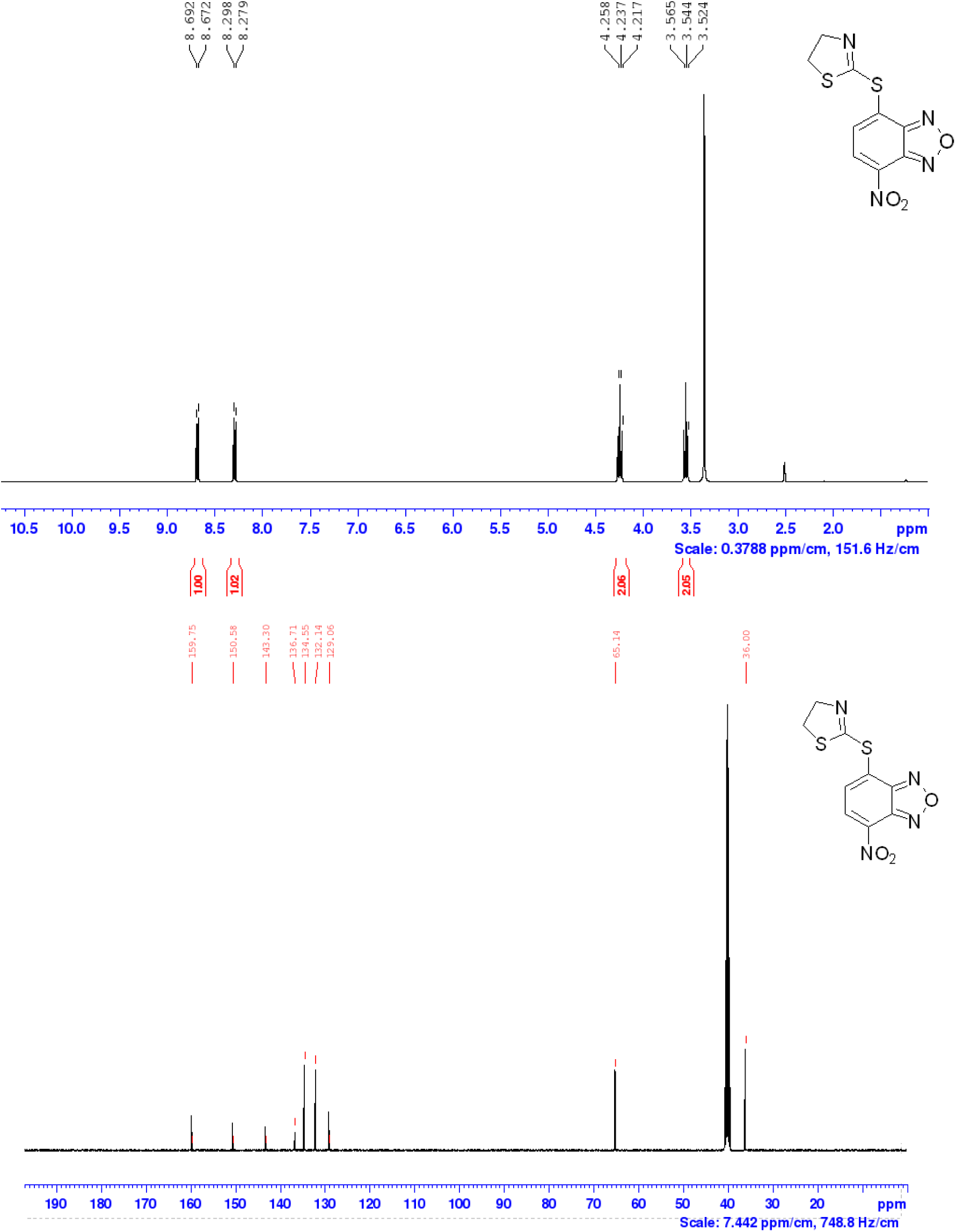

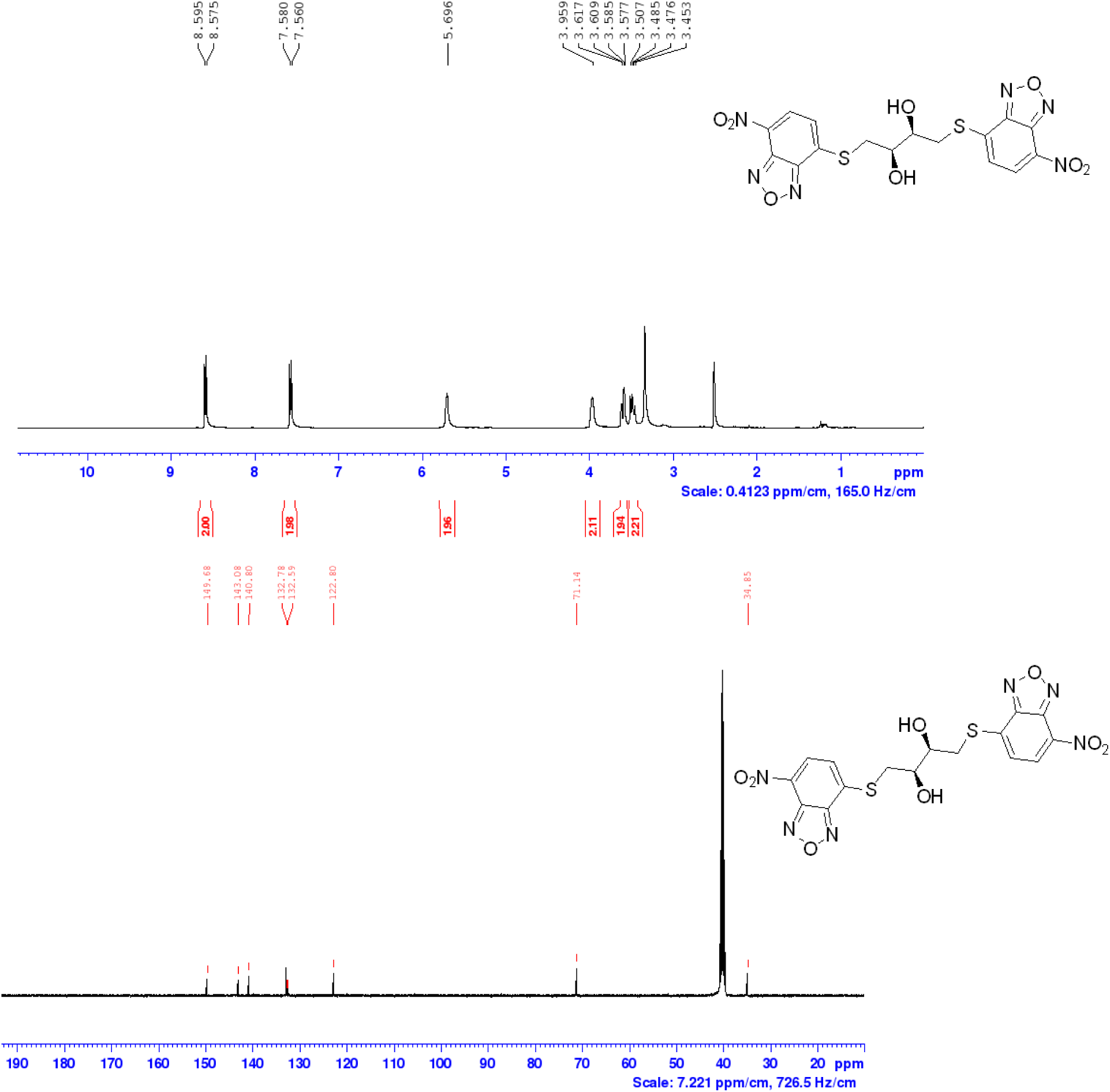

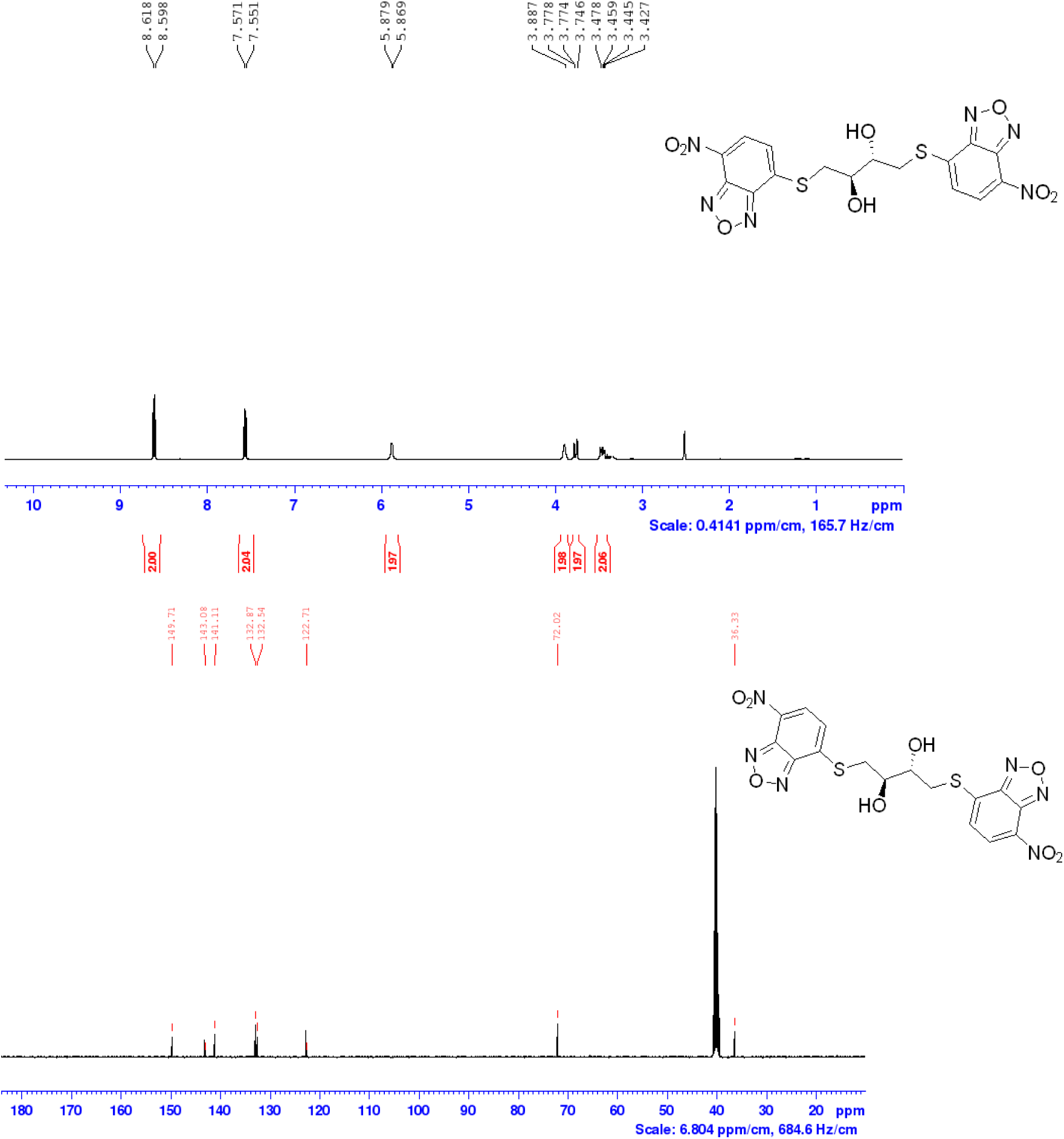

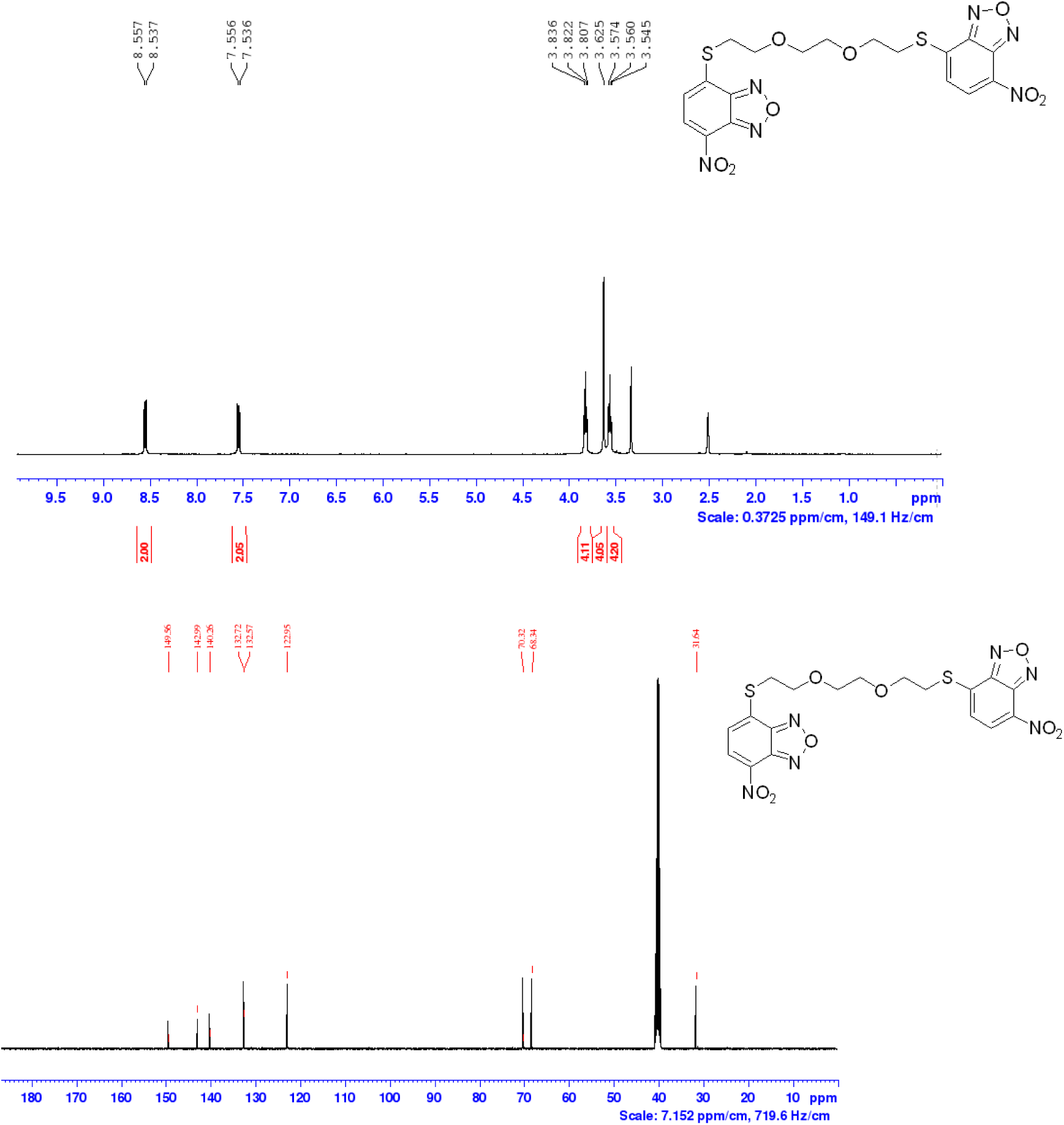

## Notes

### Competing Interest Statement

The authors have declared no competing interest.

